# Plastome convergence across heterotrophic plant lineages: genome reduction, extreme AT bias, high substitution rates, and functional persistence in the endoparasitic Mitrastemonaceae

**DOI:** 10.64898/2026.04.27.721060

**Authors:** Maria Emilia Roulet, Leonardo Gatica-Soria, Laura Evangelina Garcia, Runxian Yu, Chenyixin Wang, Renchao Zhou, M. Virginia Sanchez-Puerta

## Abstract

The loss of photosynthesis triggers extreme plastid genome (ptDNA) decay, including complete genome loss. Of the multiple transitions to heterotrophy among angiosperms, the ptDNA status remains poorly defined in lineages such as the endophytic Mitrastemonaceae (Ericales). Adopting a panplastome perspective, we characterized genomic variation across *Mitrastemon yamamotoi* individuals, assembling two complete circular ptDNAs and re-evaluating all available genomic resources for the genus. Our results reveal a highly minimized ptDNA (18–26 kb) with extreme AT content (>77%) and loss of the typical quadripartite architecture. Despite the absence of the stabilizing inverted repeats, the *Mitrastemon* panplastome exhibits remarkable structural stability and collinearity among individuals. The reduced suite of 26 genes, which includes *accD, infA, clpP, ycf1, ycf2*, and the essential tetrapyrrole precursor *trnE*`−UUC, exhibit` elevated substitution rates. Evolutionary rate analyses (dN/dS) demonstrate that the core ribosomal suite remains under strong purifying selection (ω<1), confirming the organelle’s functional status. Furthermore, transcriptomic analysis identified a nearly complete set of nuclear-encoded DNA-RRR genes, with the notable exception of the MUTS2 surveillance system. The convergent loss of these homologs in *Mitrastemon* and another holoparasitic lineage may be linked to the shared structural instability and mutational bias. Our findings demonstrate that despite extreme genome compaction, accelerated substitution rates, and severe AT-bias, the *Mitrastemon* panplastome remains quite stable, providing a definitive genomic framework for understanding plastid evolution within the endoparasitic Mitrastemonaceae.

## INTRODUCTION

The transition from autotrophy to heterotrophy involves not only physiological and morphological adaptations but also profound modifications in organellar compartments. This shift, which encompasses both mycoheterotrophic and parasitic strategies, has resulted in the independent loss of photosynthesis across numerous angiosperm lineages. Among these, parasitism represents a lifestyle dependent upon plant hosts that has arisen independently in at least 12 lineages (Cai, 2023; Nickrent, 2020). Within this spectrum of parasitism, endoparasitic plants represent the most extreme manifestation of this lifestyle (Teixeira-Costa and Suetsugu, 2023). By spending nearly their entire life cycle embedded within host tissues, endoparasites offer a unique evolutionary model to examine the functional limits and structural reconfigurations of organellar genomes, particularly the plastid genome (ptDNA).

In the ptDNA, the transition to heterotrophy is accompanied by a predictable trajectory of genomic decay. As photosynthetic constraints are relaxed, genes associated with photosystems, cytochrome complex, and ATP synthase are typically lost or pseudogenized, while a minimal suite of genes is selectively retained to support vital biosynthetic processes that remain localized within the plastid (Graham et al., 2017; Wicke and Naumann, 2018). In addition, the ptDNA undergoes convergent modifications: structural rearrangements, including the collapse of the typical quadripartite architecture, extreme AT-bias, and a significant acceleration in substitution rates (Arias-Agudelo et al., 2019; Bellot and Renner, 2016; Ceriotti et al., 2021, 2025; Ramírez-Ramírez et al., 2025; Roquet et al., 2016; Su et al., 2019; Wicke et al., 2013). In its most extreme manifestation, this degenerative process can result in the complete loss of the plastid genome, a phenomenon documented in the holoparasitic families Rafflesiaceae, Mystropetalaceae, and *Cuscuta* (Banerjee and Stefanović, 2023; Cai et al., 2021; Molina et al., 2014; Yu et al., 2025).

Comparative analyses of plastid genomes retained across diverse non-photosynthetic angiosperms have unveiled essential organellar functions that persist long after the loss of photosynthesis. Furthermore, these investigations have pushed the known boundaries of organellar evolution, revealing remarkable genomic plasticity such as record-breaking AT-bias and unprecedented genetic code modifications (Su et al., 2019). However, determining whether such extreme decay follows a universal trajectory requires the exploration of hitherto uncharacterized lineages, such as the endoparasitic Mitrastemonaceae (Ericales). Addressing this phylogenetic gap is crucial to defining the structural limits of plastome evolution and identifying the core non-photosynthetic requirements that may prevent total genomic loss in heterotrophic plants.

By adopting a panplastome perspective, we characterized the full extent of genomic variation across multiple individuals of *Mitrastemon yamamotoi*, allowing us to distinguish between accession-specific features and the strictly conserved functional core of the species. We assembled and manually curated two complete circular ptDNAs of *Mitrastemon* and performed a comprehensive re-evaluation and re-annotation of all available genomic resources for the genus to ensure a consistent comparative framework. Furthermore, we conducted phylogenomic and evolutionary rate analyses to assess the substitution rates and the selective constraints acting on the remaining gene suite.

Our results reveal a remarkable degree of structural conservation and collinearity among *Mitrastemon*, defining a stable panplastome in *M. yamamotoi*, despite extreme genome reduction and restructuring and elevated AT content and substitution rates. This study offers a genomic baseline for understanding the evolution of ptDNAs in the endoparasitic Mitrastemonaceae lineage.

## MATERIALS AND METHODS

### Sample collection and assembly of the plastid genome

A specimen of *Mitrastemon yamamotoi* #1 was sampled on October 1, 2020, in Menghai, Yunnan, China (22.0122°N, 100.3775°E), where it was found parasitizing the roots of *Castanopsis* sp. To minimize host DNA contamination, total genomic DNA was extracted exclusively from floral tissues using the CTAB protocol (Doyle, 1991). Following the construction of a 350-bp insert library, sequencing was performed on the Illumina NovaSeq 6000 (v.1.5) platform. Quality-filtered reads were subsequently archived in the NCBI Sequence Read Archive under accession SRR34031518. Additionally, we included a second dataset corresponding to a different individual of *Mitrastemon yamamotoi* #2, downloaded from the SRA (SRR16213606).

De novo plastid genome assemblies were performed independently for both *Mitrastemon* datasets using GetOrganelle v.1.7.1 (Jin et al., 2020), with default parameters for plastome assembly and k-mer sizes of 55, 73, and 89, using the embplant_pt plastid reference database. Resulting contigs were inspected in Bandage v.0.8.1 (Wick et al., 2015), and the contigs identified as being of plastid origin were identified via BLASTn searches against plastid genomes of Ericales retrieved from GenBank (Table S1). Plastid-derived contigs were manually curated, edited, and joined in Consed v.29 (Gordon and Green, 2013) to reconstruct complete circular genomes. Final assemblies were validated by read mapping in Consed to ensure consistent paired-end coverage across junctions. To estimate DNA read depth across the ptDNAs, paired-end read mapping was performed in Bowtie2 v.2.4.4 (*parameters*: -I 100 X 350 --end-to-end --very-sensitive --no-contain --no-discordant --no-mixed) (Langmead and Salzberg, 2012).

In addition to these assemblies, we included two unpublished plastid genomes from *Mitrastemon* available in Genbank (NCBI) for comparative analyses: *M. yamamotoi* #3 (GenBank accession NC_080971) and *M. yamamotoi var. kanehirai* (GenBank accession MF372930).

### Plastid genome characterization

The plastid genomes of *Mitrastemon* were annotated using GeSeq (Tillich et al., 2017) with a comprehensive combination of BLAT, HMMER, and third-party tools. For the BLAT search, protein identity was set to 40% and rRNA/tRNA/DNA identity to 85%, enabling the annotation of CDS, tRNA, and rRNA genes. The HMMER profile search was performed using the “Chloroplast land plants” model, with annotation of both CDS and rRNA features. Additional tRNA prediction was conducted using ARAGORN v.1.2.38 (with the Bacterial/Plant Chloroplast genetic code, intron length max = 3000 bp), tRNAscan-SE v.2.0.7 (organellar tRNA mode), and Chloë v.0.1.0. Default parameters were used unless otherwise specified. Final annotations were manually curated in Geneious Prime® 2024.0.4 (Kearse et al., 2012). The annotated plastid genomes of *M. yamamotoi* #1 and *M. yamamotoi* #2 were deposited in NCBI (Genbank accession PX991927) or available in Figshare (Roulet et al., 2026), respectively.

To ensure consistency across datasets, we re-annotated the available plastid genomes of *M. yamamotoi #3* and *M. yamamotoi var. kanehirai* using the same GeSeq parameters and manual curation workflow. Repetitive sequences in the plastid genomes of the four individuals of *Mitrastemon* were identified using the get_repeats.sh script (Gandini et al., 2019) with default parameters, except for the -perc_identity flag set to 90, which defines repeats as sequences sharing at least 90% identity.

To estimate RNA read depth across the assembled plastid genome and its genes in *M. yamamotoi* #1, we downloaded publicly available RNA-seq data for *Mitrastemon yamamotoi* (SRR28027597) (Carruthers et al., 2024) and confirmed its unstranded library format using Salmon’s (v.1.20) automatic detection (-l A). We used the plastid genome assembly and the coding sequences (CDS) of each gene, extended by 300 bp at both ends to improve read mapping at gene boundaries. RNA reads were aligned using Bowtie2 (Langmead and Salzberg, 2012) with the “end-to-end” and “very-sensitive” presets, along with the --no-discordant and -R 10 options.

### Phylogenetic analyses

A phylogenetic analysis was conducted using a concatenated dataset of 14 plastid protein-coding genes (*clpP, rpl2, rpl16, rpl36, rps2, rps3, rps4, rps7, rps8, rps11, rps12, rps14, rps18*, and *rps19*) obtained from *Mitrastemon* individuals and other angiosperm species (Table S1). Each gene was aligned independently using MAFFT v.7.487 with the --localpair and --maxiterate 1000 parameters (Katoh and Standley, 2013). The resulting alignments were concatenated and used to infer a maximum likelihood phylogeny with IQ-TREE v.2.2.0 (Minh et al., 2020). Model selection was performed using ModelFinder (-m MFP), which identified GTR+F+R4 as the best-fit nucleotide substitution model according to the Bayesian Information Criterion (BIC). Branch support was assessed using 1000 ultrafast bootstrap replicates. The tree was visualized in FigTree v.1.4.4 and rooted according to the Angiosperm Phylogeny Group (APG) classification (The Angiosperm Phylogeny Group et al., 2016).

### Evolutionary rate analyses

To evaluate evolutionary rates in plastid protein-coding genes of *Mitrastemon*, we selected 13 genes commonly retained across *Mitrastemon* and autotrophic angiosperms (*clpP, rpl2, rpl16, rps2, rps3, rps4, rps7, rps8, rps11, rps12, rps14, rps18*, and *rps19*). Evolutionary rate estimates (non-synonymous [dN] or synonymous [dS] substitutions per non-synonymous or synonymous site, respectively, and dN/dS [ω]) were calculated using the CodMLl program from the PAML v.4.7 package (Yang, 2007). We used the branch model (model = 1) to allow variation of ω among branches, with the F3×4 codon frequency model (CodonFreq = 2) and seqtype = 1 for codon data. The transition/transversion ratio was set to kappa = 2, and fix_omega = 0 allowed ω to be estimated. The input phylogenetic tree was constructed according to the most recent APG (2016) classification and the phylogeny obtained in this study. To process the output of CodeML, we used a custom shell pipeline and an R script, available in Figshare (Roulet et al., 2026), that extracted terminal branch rates (dN, dS, and ω), parse labeled trees and visualize root-to-tip distances.

### Identification of plastid DNA-RRR and plastid ribosomal protein encoding genes in the *Mitrastemon* transcriptome

To identify nuclear-encoded plastid DNA recombination, repair, and replication (DNA-RRR) genes and plastid ribosomal protein encoding genes, we used the publicly available *Mitrastemon* RNA-seq data (SRR28027597) (Carruthers et al., 2024). The transcriptome was *de novo* assembled using Trinity v.2.15.0 with default parameters, except for a minimum contig length of 100 bp (--min_contig_length 100) (Grabherr et al., 2011). Open Reading Frames (ORFs) were predicted with TransDecoder, specifying a minimum length of 100 amino acids (TransDecoder.LongOrfs -t Trinity_out.Trinity.fasta -m 100).

DNA sequences of 30 plastid DNA-RRR genes from *Arabidopsis thaliana* (Ceriotti et al., 2022; Gualberto and Newton, 2017; Zhang et al., 2016) were used as queries in BLASTN and TBLASTX searches against the assembled *Mitrastemon* transcriptome and the predicted ORFs, respectively. Candidate ORFs were validated as homologs of the corresponding *Arabidopsis* genes through reciprocal BLAST searches. Similarly, sequences of nuclear-encoded plastid ribosomal genes from *Arabidopsis* (Scarpin et al., 2023) were used to search for homologous transcripts in the *Mitrastemon* transcriptome and ORFs.

## RESULTS

### Architecture, AT-bias, and repeat landscape of the minimized *Mitrastemon* plastid genome

The ptDNAs of *M. yamamotoi* #1 and #2 assembled as circular molecules with lengths of 25,623 bp and 25,999 bp, respectively, supported by gap-free read coverage across their entire length (Figure S1A). A representative ptDNA map for *M. yamamotoi* is shown in Figure 1. To further explore plastome evolution across individuals, we compared these new assemblies with sequences from *M. yamamotoi* #3 and *M. yamamotoi* var. *kanehirai* available in Genbank. Our comparative analysis revealed length variation, ranging from 18,252 bp to 25,999 bp (Figure S2). The variation in ptDNA length among individuals is mainly attributed to differences in intergenic regions due to the accumulation of indels. *Mitrastemon* plastid genomes are highly AT-rich, with values exceeding 77%, and lack the typical quadripartite genome architecture. The repetitive content of the ptDNA of the four individuals of *Mitrastemon* was similar among them, ranging from 4.56% to 6.62%. In *M. yamamotoi* #1, we identified 36 tandem repeats spanning 1,494 bp (5.83%) and a single pair of short, dispersed repeats (<100 bp), for a total repeat content of 1,519 bp (5.93%). In *M. yamamotoi* #2, 45 tandem repeats were found (1,533 bp, 5.90%), along with six pairs of short, dispersed repeats, summing to 1,722 bp (6.62%) of repetitive content (Table 1, Table S2).

**Table 1.**
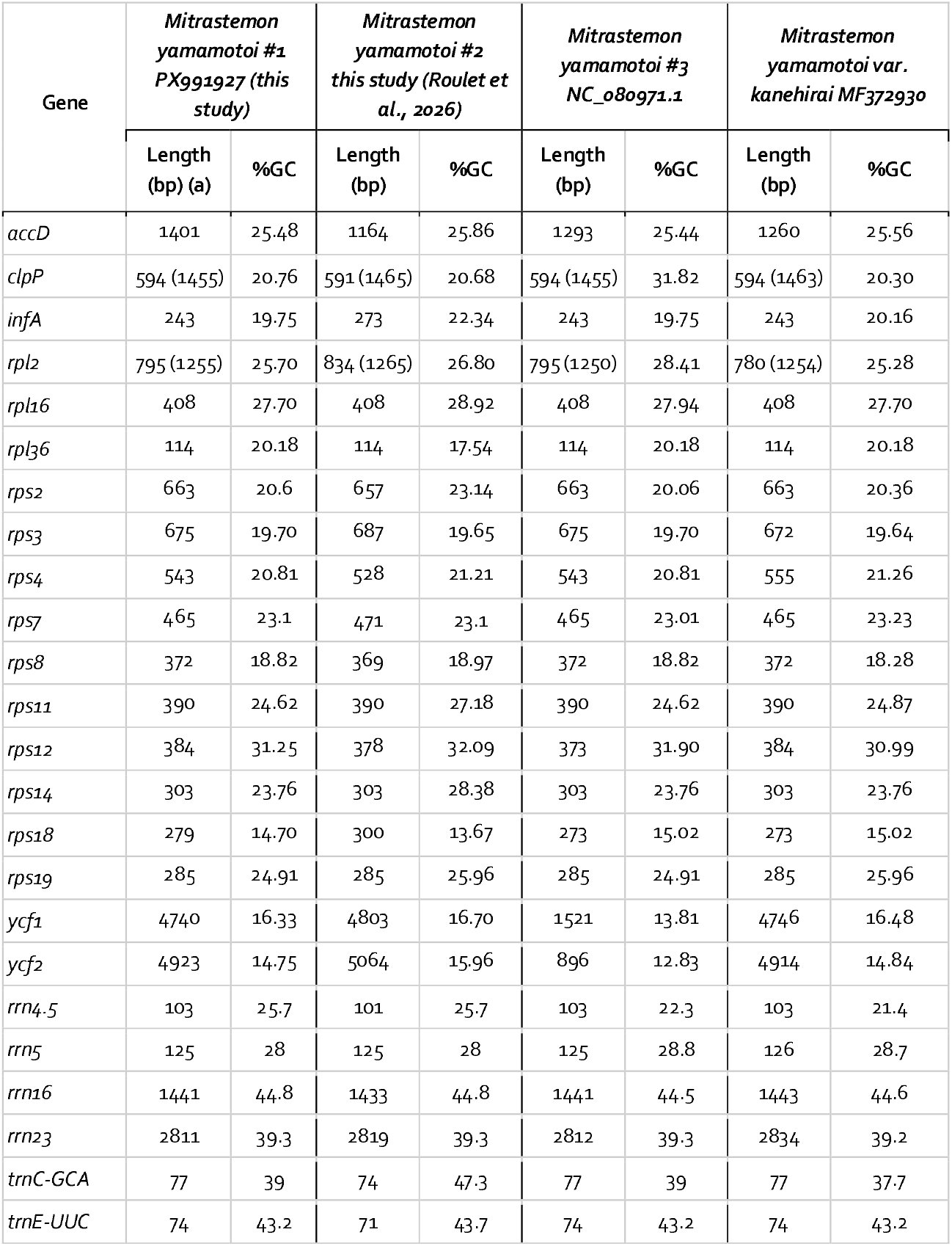
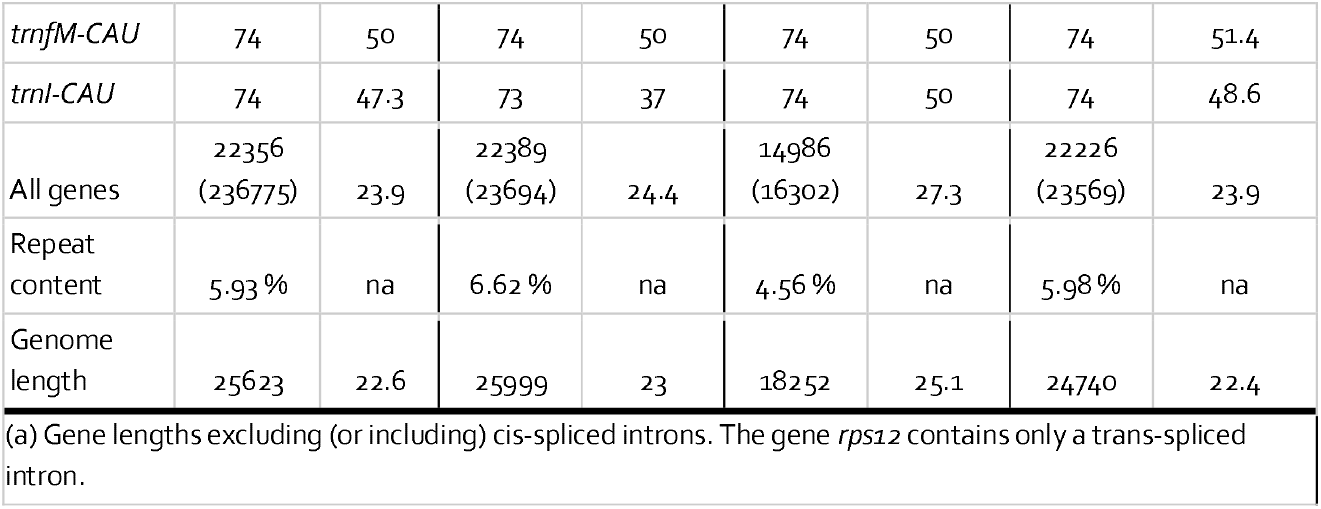
Features of *Mitrastemon* plastid genes.

**Figure 1.**
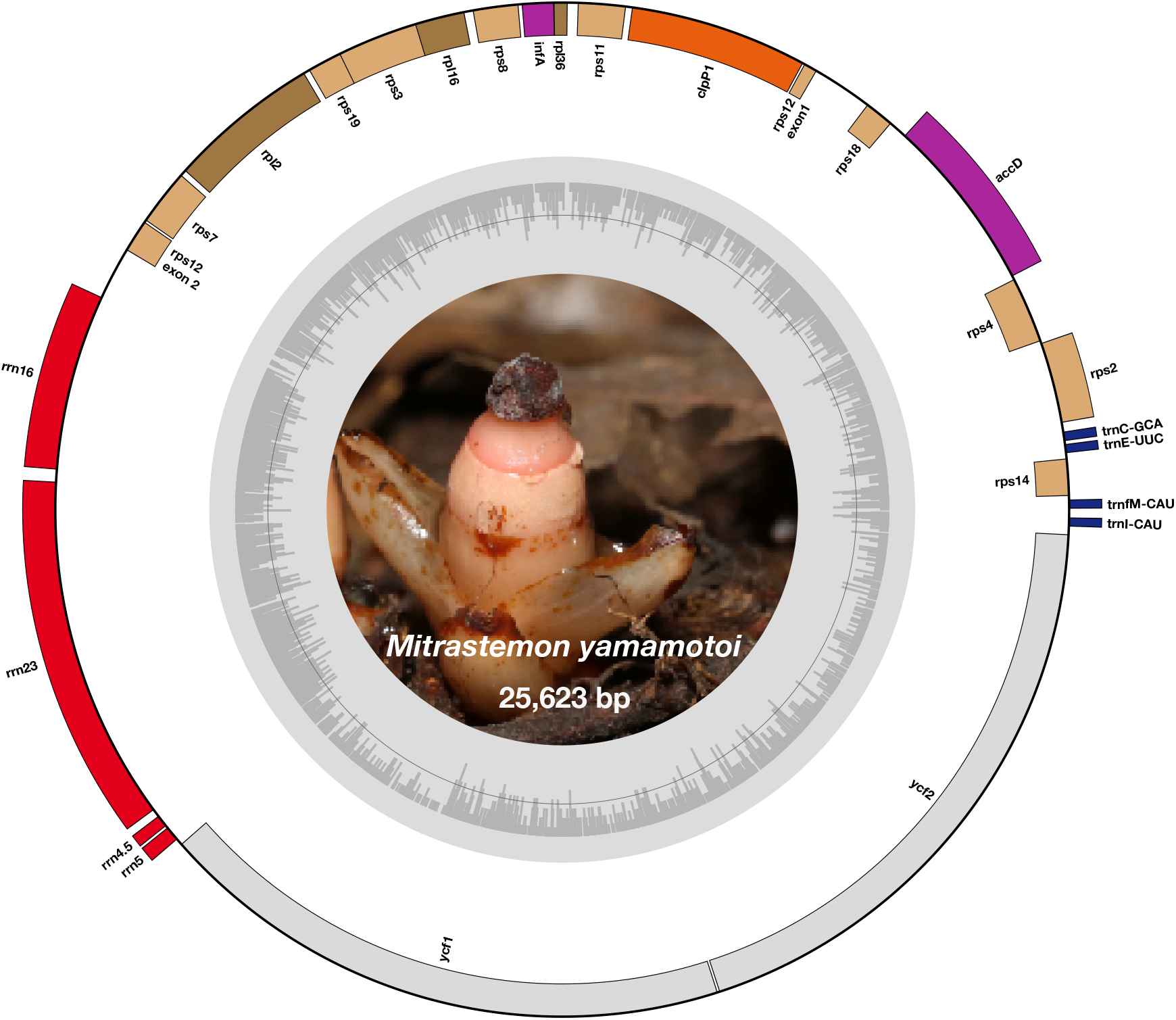
Circular map of the *Mitrastemon yamamotoi* plastid genome. The plastid genome (ptDNA) of *M. yamamotoi* individual (#1) is 25,623 bp in length. Genes transcribed clockwise and counter-clockwise are shown on the outer and inner sides of the circle, respectively. The inner histogram represents GC content, with an average of 22.6%. The central image shows the flower of *M. yamamotoi* (photo by Runxian Yu).

### Reduced plastid gene content in *Mitrastemon*

The *Mitrastemon* panplastome lacks genes involved in photosynthesis, ATP synthase, RNA polymerase, and splicing factors, resulting in a shared set of 26 genes: 18 protein-coding, four rRNA, and four tRNA genes (Table 1). These genes occupy 82–87% of the total plastome length (Table 1). *Mitrastemon* individuals retain a minimal set of four tRNA genes: *trnC-GCA, trnE-UUC, trnfM-CAU*, and *trnI-CAU*. Most genes are intronless; however, *clpP* and *rpl2* retain cis-spliced introns, and *rps12* includes a trans-spliced intron in all individuals. Stop codons were annotated for all protein-coding genes, with TAA being the most frequent, followed by TAG and TGA (Table S3). A comparative analysis of the four *Mitrastemon* plastomes shows identical gene collinearity (Figure S2), and no structural rearrangements, inversions, or translocations, even in the absence of the stabilizing inverted repeats (Maréchal and Brisson, 2010). However, gene order is not conserved relative to the photosynthetic species *Barringtonia asiatica* (Ericales; NC_070213.1).

Besides, *Mitrastemon* ptDNAs exhibit pronounced reduction in gene lengths, especially in large protein-coding genes. For example, in *Mitrastemon clpP* spans 1,455–1,465 bp, and the longest *ycf1* and *ycf2* sequences reach 4,803 bp and 5,064 bp, respectively (Table S3), compared with the photosynthetic *Barringtonia asiatica* (Ericales; NC_070213.1) exhibiting a *clpP* of 2,248 bp, a *ycf1* of 5,676 bp, and *ycf2* of 6,918 bp. These reductions reflect the trend of gene compaction associated with parasitism.

Further signs of ptDNA compaction in *Mitrastemon* are observed in the presence of overlapping genes. All four *Mitrastemon* ptDNAs exhibit consistent overlaps within the cluster *rpl16–rps3–rps19*, three of them within the cluster *rpl36–infA*, with an additional *infA–rps8* overlap detected in *M. yamamotoi* #2 (Figure S2).

We reconstructed a maximum likelihood phylogeny based on a concatenated alignment of 14 conserved plastid genes (6,825 bp total; Figure S3). All *Mitrastemon* accessions formed a strongly supported clade (bootstrap support = 100%), confirming their monophyly. Moreover, *M. yamamotoi* #2 forms a sister lineage to a clade containing *M. yamamotoi* #1, #3, and *M. yamamotoi* var. *kanehirai*.

### Comparative plastid genome features across non-photosynthetic and autotrophic angiosperms

We compared the plastid gene content of *Mitrastemon* with that of the other endoparasitic angiosperms available, namely *Cytinus* (Cytinaceae), *Apodanthes*, and *Pilostyles* (Apodanthaceae) (Table S3). Despite their independent evolutionary origins, all three lineages exhibit highly reduced plastid gene content, though with distinct patterns and degrees of gene loss. Furthermore, an analysis of ptDNA size and AT content across a broader range of angiosperms, encompassing holoparasites, mycoheterotrophs, endoparasites, and photosynthetic representatives, reveals a negative correlation between genome length and AT content across the ptDNAs (Figure 2, Table S1). While photosynthetic taxa cluster at lengths of 120–160 kb with 61–64% AT content, heterotrophic species exhibit a drastic shift toward minimized, AT-richer ptDNAs (>70% AT). *Mitrastemon* exceeds the AT levels of other endoparasites like *Cytinus hypocistis* (70.8% AT) (Roquet et al., 2016) and is comparable to the endoparasite *Pilostyles aethiopica* (75.8% AT) (Bellot and Renner, 2016), but remains below the record-setting values of *Balanophora spp*. (holoparasite), which can reach an extraordinary 88.4% AT (Chen et al., 2020; Su et al., 2019; Yu et al., 2025).

**Figure 2.**
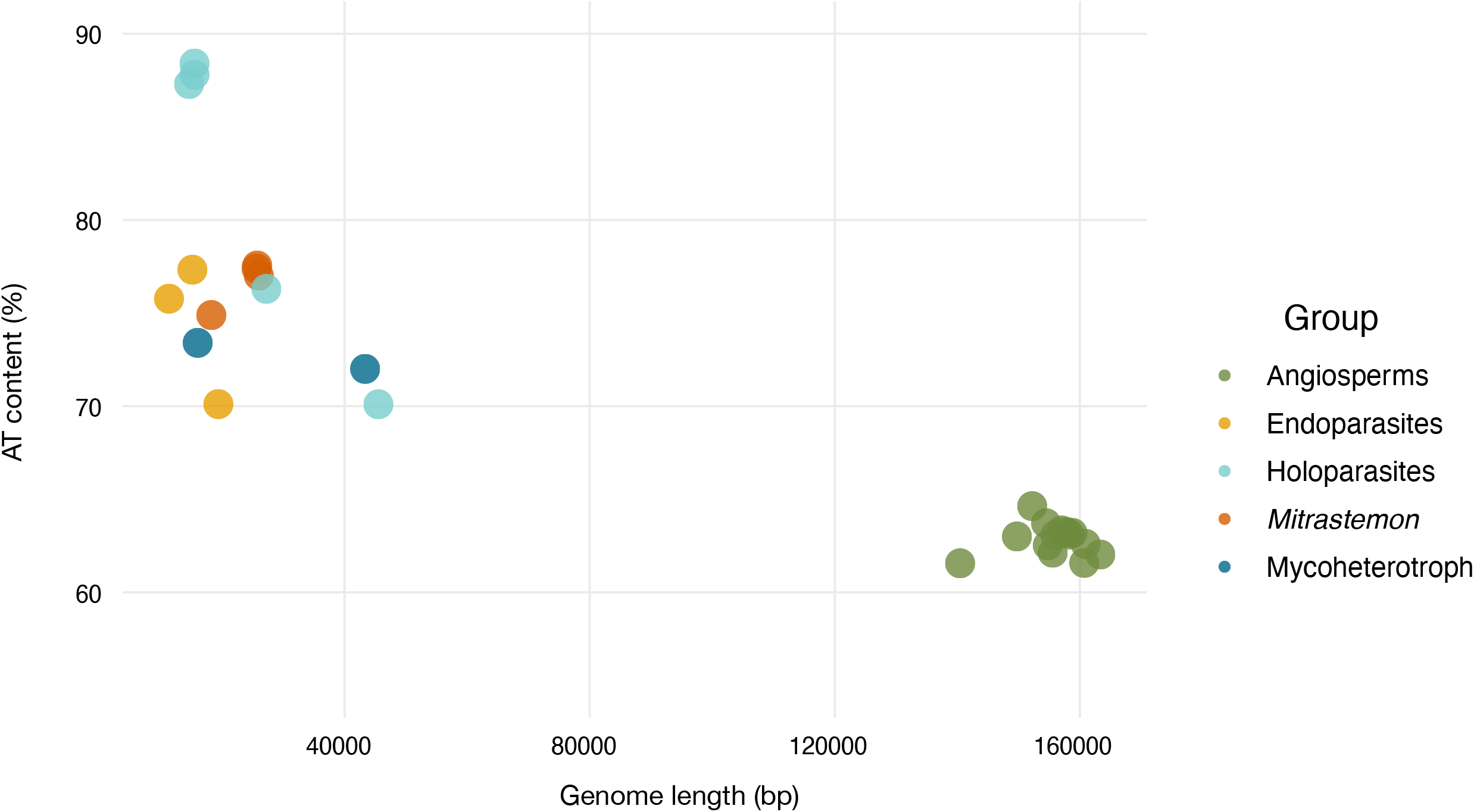
Relationship between plastid genome size and AT content across representative angiosperms and holoparasitic species. The bubble plot shows genome size (x-axis) and AT content (%) (y-axis) of complete plastid genomes. Species are colored by phylogenetic or ecological group: Holoparasites (light-blue), Mycoheterotroph (blue), Endoparasites (yellow), *Mitrastemon* (orange), and other autotrophic angiosperms (green).

### Elevated substitution rates in *Mitrastemon* plastid genes

Previous studies have consistently shown that parasitic plants, both holo- and endoparasites, exhibit accelerated rates of molecular evolution compared to their non-parasitic relatives (Bromham et al., 2013; Ceriotti et al., 2022; Nickrent and Starr, 1994; Roquet et al., 2016; Su et al., 2019; Yu et al., 2025). Following the hypothesis that the endoparasitic *Mitrastemon* lineage undergoes similar evolutionary shifts, we calculated root-to-tip substitution rates (dN and dS) for 13 conserved plastid genes using branch models in CodeML (Table S4, Figure 3 and S4). Both nonsynonymous (dN) and synonymous (dS) substitution rates were markedly elevated in *Mitrastemon* compared to photosynthetic angiosperms (including close relatives within the Ericales), with mean root-to-tip dN and dS values of 0.297 and 2.198, respectively (Table S4).

**Figure 3.**
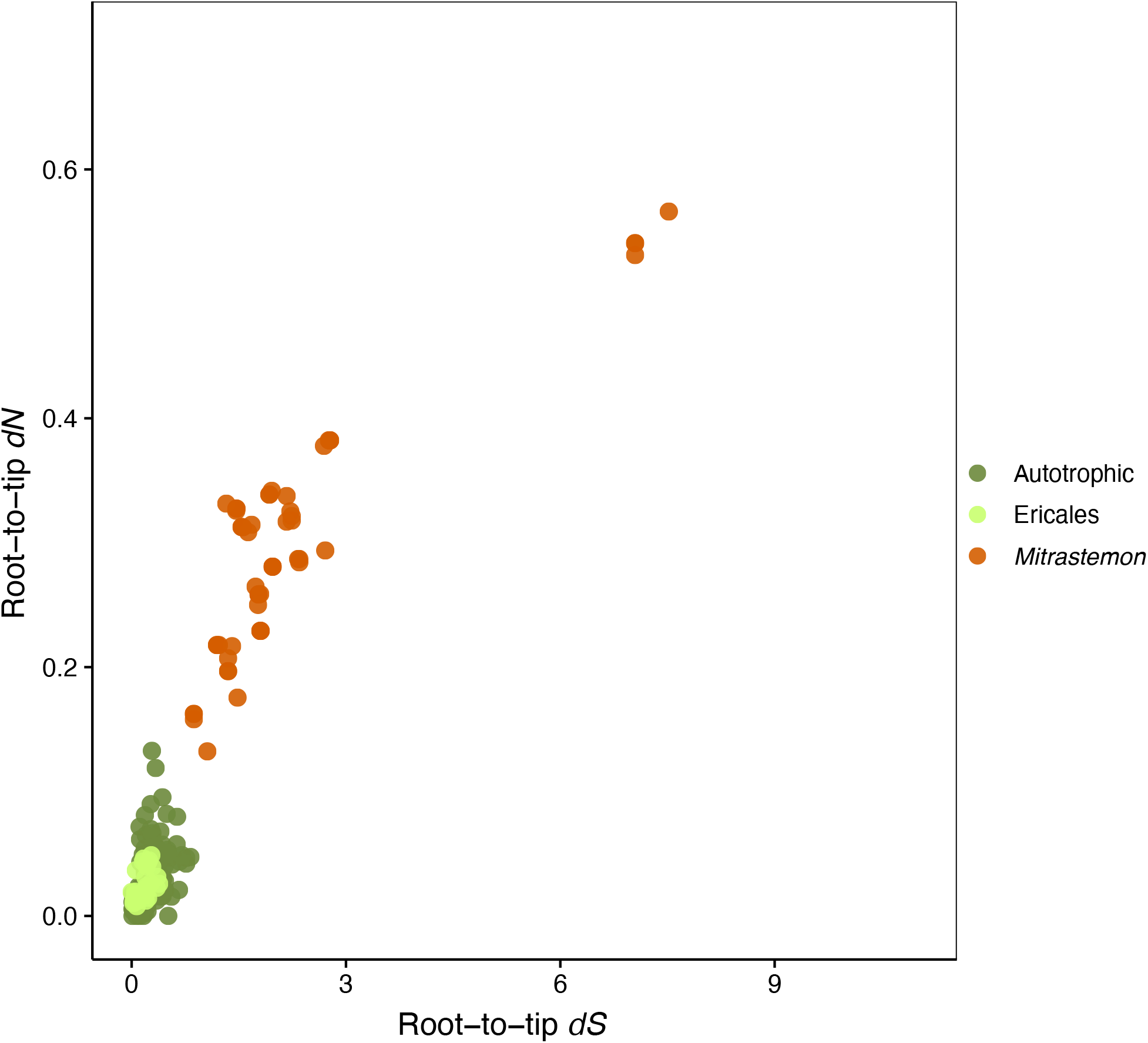
Root-to-tip substitution rates for plastid genes. Each point represents the cumulative dN and dS values for each plastid gene and species, obtained from codeml branch models. Species are color-coded by group: *Mitrastemon*, Ericales, and autotrophic angiosperms.

Substitution rates showed some degree of heterogeneity both within *Mitrastemon* accessions and across genes, with several genes exhibiting particularly high dN or dS values relative to autotrophic species (Figure 3 and S4). Individual gene trees used for rate estimation are shown in Figure S5.

Based on the ω (dN/dS) estimates across plastid genes in *Mitrastemon*, all retained genes appear to be evolving under purifying selection (ω < 1), indicating functional constraint in these retained genes (Table S4). Besides, functional confirmation through mapping of RNA-seq data determined the transcription and splicing of these genes in the plastid genome of *Mitrastemon* (Figure S1B).

### Conserved nuclear genes for plastid ribosomes in *Mitrastemon*

To evaluate whether the plastid ribosome remains functionally complete despite plastid gene losses and elevated substitution rates in plastid genes, we surveyed the presence of nuclear- and plastid-encoded ribosomal subunits in *Mitrastemon*. We searched for the 33 plastid ribosomal proteins encoded in the nuclear genome of *Arabidopsis* (Scarpin et al., 2023) and identified transcripts corresponding to all 33 subunits in the *Mitrastemon* transcriptome (Table S5).

In addition, 13 ribosomal protein genes were identified in the *Mitrastemon* plastid genome (Table 1 and Figure S2). However, eight plastid ribosomal genes present in *Arabidopsis* ptDNA are absent from the *Mitrastemon* plastome (Table S5). Of these eight missing plastid-encoded genes, three were recovered in the *Mitrastemon* transcriptome, indicating that they are likely nuclear-encoded in this lineage (Table S5). Overall, plastid ribosomal subunits are largely complete in *Mitrastemon*.

### Plastid-targeted DNA-RRR proteins in *Mitrastemon*

We examined the transcriptome of *Mitrastemon* to assess the presence of genes involved in DNA-RRR that might explain the structural re-arrangements and elevated substitution rates and AT content of the plastomes in *Mitrastemon*. We identified almost all 30 nuclear-encoded DNA-RRR genes whose products are targeted to the plastid in *Arabidopsis* in the transcriptome of *M. yamamotoi* (Table 2). The only two missing genes were the MutS homologs MUTS2-1 and MUTS2-2, which are absent or not expressed. A previous assessment of the completeness of the assembled transcriptome showed that the transcriptome assembly of *Mistrastemon* may be considered fairly complete for an endoparasitic plant (Roulet et al., 2025). This suggests that the missing genes are not a result of an incomplete transcriptome, but the lack of expression of both MUTS2-1 and MUTS2-2 genes might result from the loss of those genes from the nuclear genome of *Mitrastemon*.

**Table 2.**
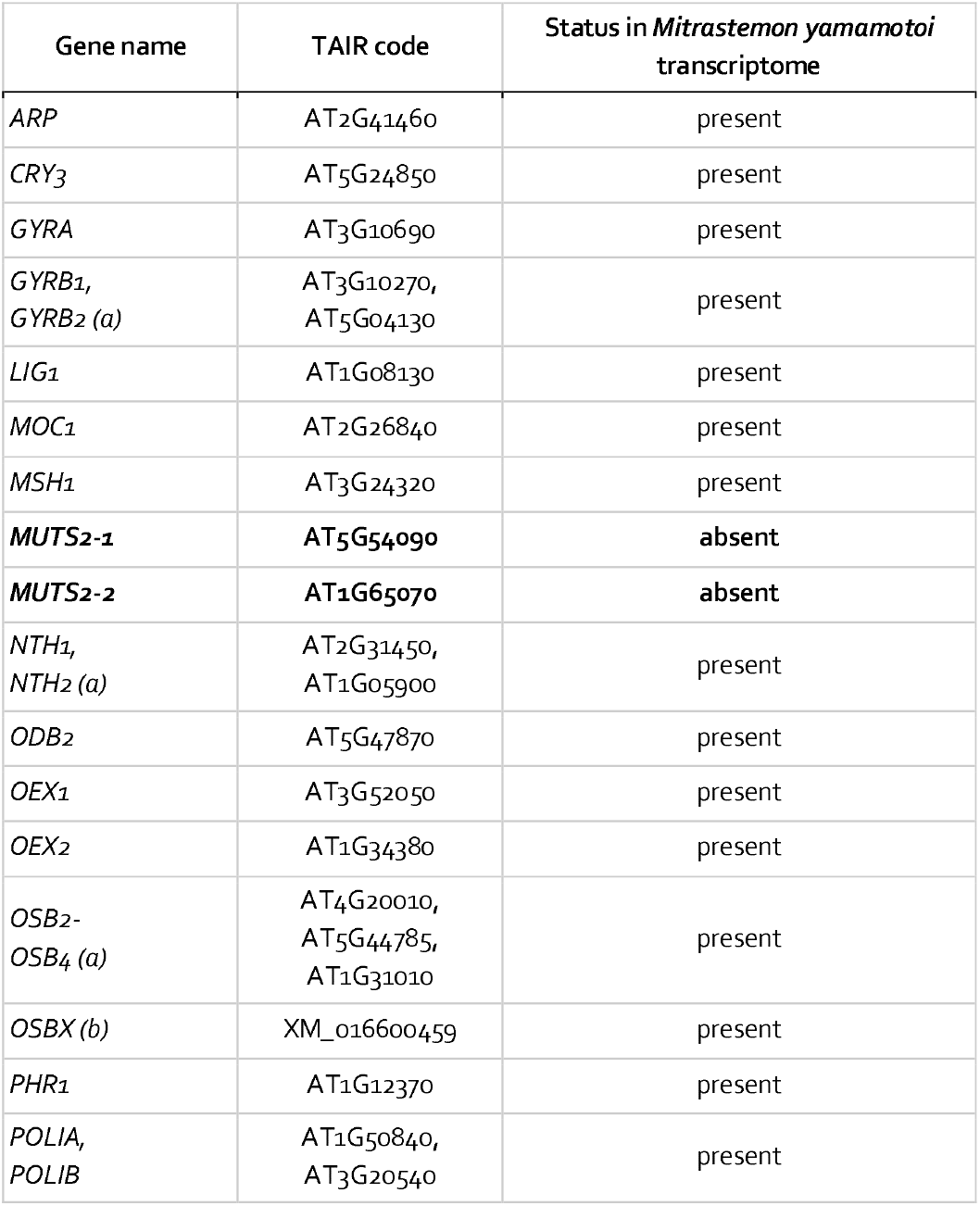
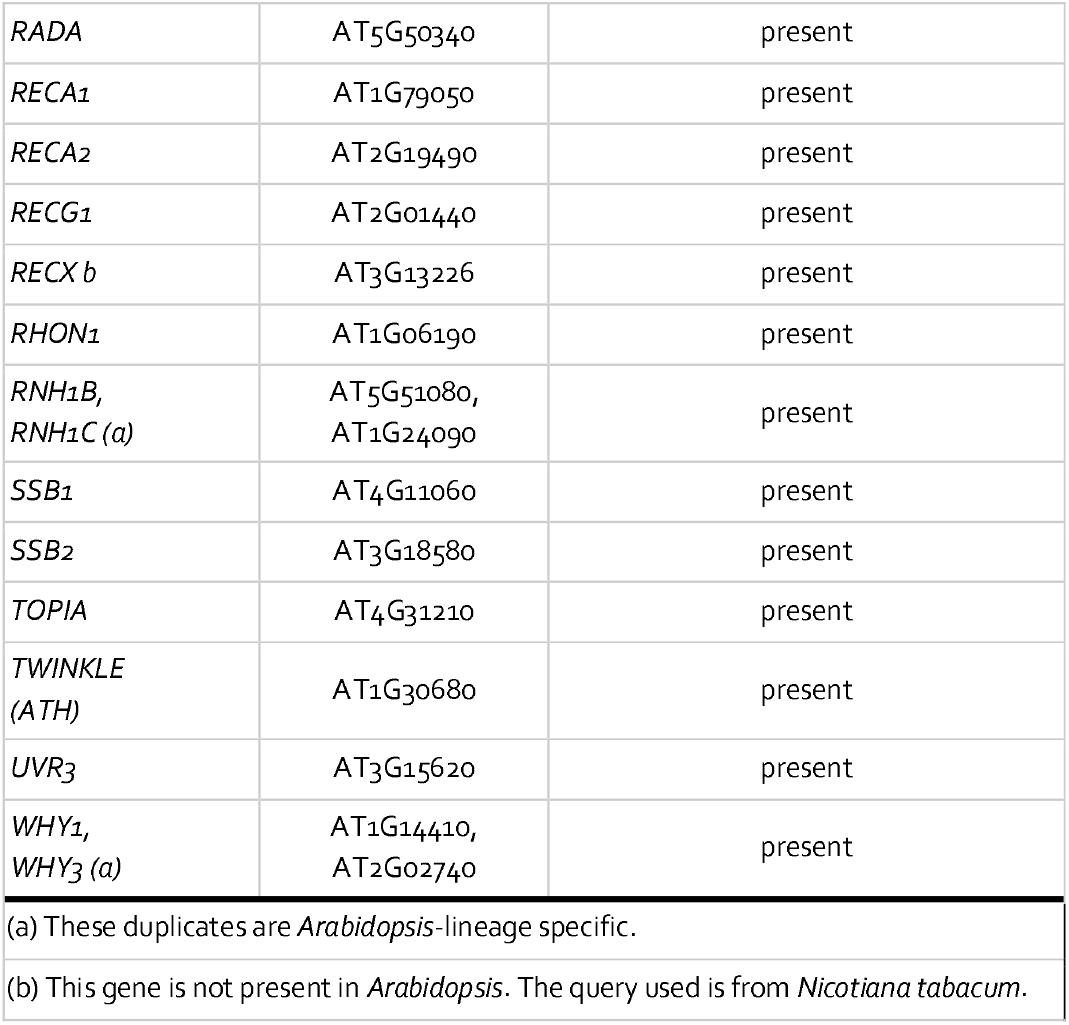
Identification of plastid DNA-RRR genes in the transcriptome of *Mitrastemon yamamotoi*.

## DISCUSSION

### Panplastome architecture in *Mitrastemon*

Our study provides the first comparative view of complete plastomes in the endoparasitic family *Mitrastemonaceae*. The *Mitrastemon* panplastome is highly reduced (∼26 kb), retaining only 18 protein-coding genes, along with four rRNAs and four tRNAs (Table 1, Figure S2). No genes related to photosynthesis, RNA polymerase, or the ATP synthase complex remain. This reduced gene complement is consistent with patterns observed in other endoparasitic lineages and exemplifies the convergent trajectory of plastome reduction following photosynthesis loss (Arias-Agudelo et al., 2019; Bellot and Renner, 2016; Ramírez-Ramírez et al., 2025; Roquet et al., 2016).

This panplastome is highly stable, with identical gene content and order among individuals, despite massive genome reduction and length variation. Overlapping gene arrangements (e.g., *rpl16–rps3–rps19*), along with severely shortened intergenic regions, point to strong mechanisms driving genome compaction, features also observed in other highly reduced plastomes such as those of Balanophoraceae and Apodanthaceae (Arias-Agudelo et al., 2019; Ceriotti et al., 2021, 2025; Ramírez-Ramírez et al., 2025; Su et al., 2019; Yu et al., 2025). Notably, even in the absence of the inverted repeats, which typically stabilizes the genome (Maréchal and Brisson, 2010), no other structural rearrangements, inversions, or translocations were detected in *Mitrastemon* ptDNAs. The high coding density and frequent gene overlaps observed in *Mitrastemon* may impose a physical constraint on genomic rearrangements (Bellot and Renner, 2016; Wicke et al., 2013).

### Retention of essential metabolic functions in the reduced plastid genome

Beyond structural stasis across individuals, the *Mitrastemon* panplastome retains a minimal but evolutionary relevant gene set. The retention of a core set of ribosomal protein genes (e.g. *rpl2, rps3, rps4*, and *rps12*) in *Mitrastemon* aligns with observations in other endoparasitic lineages, including Cytinaceae and Apodanthaceae (Arias-Agudelo et al., 2019; Bellot and Renner, 2016; Ramírez-Ramírez et al., 2025; Roquet et al., 2016). Particularly notable is the retention of *accD, clpP, ycf1*, and *ycf2*, often considered the “last bastions” of the plastid genome prior to total loss (Wicke et al., 2013; Wicke and Naumann, 2018). The persistence of *accD* and *clpP* in all endoparasitic species suggests that the plastid continues to fulfill essential metabolic roles, particularly in fatty acid biosynthesis and intra-plastid proteolysis. Furthermore, the conservation of this reduced but putatively functional gene set in *Mitrastemon* parallels patterns seen in other holoparasitic plants, such as *Hydnora visseri* (Hydnoraceae) and members of the Balanophoraceae (Ceriotti et al., 2021, 2025; Naumann et al., 2016; Su et al., 2019). Interestingly, this pattern of extreme reduction is also observed within the order Ericales in the mycoheterotrophic species *Monotropa uniflora* (Ericaceae). Despite their different heterotrophic strategies, both *Mitrastemon* and *Monotropa* exhibit remarkably similar plastome sizes (∼26 kb) and retain a shared suite of housekeeping and metabolic genes, including *accD, infA*, and *clpP* (Liu et al., 2020).

A striking feature in the *Mitrastemon* panplastome is the retention of the `tr`*□□*− `UUC` gene, highlighting the persistent metabolic relevance of the plastid despite the complete loss of photosynthesis. Unlike other tRNAs commonly lost during plastome erosion, `tr`*□□* − `UUC` fulfills a dual essential role. In addition to its canonical function in translation, it serves as the initiator molecule in the five-carbon (C5) pathway for tetrapyrrole biosynthesis (Barbrook et al., 2006). The presence of `tr`*□□* − `UUC` in *Mitrastemon* and in *Cytinus* contrasts with the documented absence in other holo- and endoparasitic lineages such as *Pilostyles, Apodanthes, Lophophytum*, and *Ombrophytum* (Arias-Agudelo et al., 2019; Bellot and Renner, 2016; Ceriotti et al., 2021, 2025; Ramírez-Ramírez et al., 2025; Roquet et al., 2016; Sanchez-Puerta et al., 2023; Su et al., 2019).

### Extreme AT bias in *Mitrastemon* plastid genomes

The transition to a non-photosynthetic lifestyle in *Mitrastemon* is accompanied by severe sequence compaction and a strong bias toward AT nucleotides. Our results reveal a clear negative correlation between ptDNA length and AT content (Figure 2), a pattern that appears to be a convergent hallmark of plastid evolution in heterotrophic lineages (Arias-Agudelo et al., 2019; Bellot and Renner, 2016; Ceriotti et al., 2021, 2025; Ramírez-Ramírez et al., 2025; Roquet et al., 2016; Su et al., 2019).

*Mitrastemon* exhibits an AT content exceeding 77%. This value significantly surpasses those of other members of the order Ericales, such as the mycoheterotroph *Monotropa uniflora* (72.9%), and other endoparasites like *Cytinus hypocistis* (70.1%) (Roquet et al., 2016). While *Mitrastemon*’s AT levels are comparable to those of *Pilostyles* species (75.8%–77.3%) (Bellot and Renner, 2016), they remain below the extreme threshold observed in the holoparasite *Balanophora* spp., where AT content reaches an unprecedented 88.4% (Su et al., 2019; Yu et al., 2025). An increase in plastome AT content has consistently been shown to accompany the transition to heterotrophy, supporting a link between AT bias and the relaxation of selective constraints following the loss of photosynthetic function (Ceriotti et al., 2021, 2025; Roquet et al., 2016; Su et al., 2019; Wicke et al., 2013; Yu et al., 2025).

A compelling point of discussion is whether this extreme AT bias has led to a redefinition of the genetic code, as seen in several Balanophoraceae species that exhibit a reassignment of the TAG/TGA stop codon to encode Tryptophan (Ceriotti et al., 2021; Su et al., 2019). Despite its elevated AT content, *Mitrastemon* retains the canonical stop codon usage. Our annotation confirms that all three canonical stop codons (TAA, TAG, and TGA) remain functional, with TAA being the most frequent.

### Selective pressures on plastid coding regions of *Mitrastemon*

The plastid coding regions of *Mitrastemon* exhibit markedly elevated root-to-tip nonsynonymous (dN) and synonymous (dS) substitution rates (mean dN=0.297, dS=2.198) compared to photosynthetic relatives (Table S4). Our findings align with the trends observed in *Cytinus hypocistis*, where rates are an order of magnitude higher than in autotrophic Malvales (Roquet et al., 2016), and the holoparasite *Balanophora*, where dS values often exceed 1.5, indicating mutational saturation (Su et al., 2019). Additionally, these values are substantially higher than those typically observed in photosynthetic angiosperms, consistent with the relaxed selective constraints following the loss of photosynthesis (Wicke and Naumann, 2018).

Despite the overall acceleration, all plastid genes of the *Mitrastemon* panplastome appear to be under purifying selection, with ω values below 1. This selective constraint is an indicator that these genes remain functionally essential for plastid protein translation. A similar pattern is found in *Balanophora*, where 13 out of 14 genes show ω ≤ 0.40, and in *Cytinus*, where eight genes like *rps12* and *rpl2* exhibit lower ω values (Roquet et al., 2016; Su et al., 2019).

The selective maintenance provides evidence that the *Mitrastemon* ptDNA is not a mere vestigial remnant in transit toward total loss; rather, it remains a metabolically indispensable compartment whose protein synthesis capacity is still vital to the heterotrophic lifestyle.

### Convergent loss of the MUTS2 surveillance system in holoparasitic lineages

RRR proteins function as genomic surveillance systems that limit illegitimate recombination and promote accurate DNA repair. Their importance is underscored by the frequent absence or malfunction of these genes in lineages and mutant backgrounds characterized by extensive structural rearrangements and elevated mutation rates (Odahara et al., 2015; Wu et al., 2020; Yu et al., 2025; Zhang et al., 2016). In this context, the apparent absence of the two plastid-targeted MUTS2 homologs (MUTS2-1 and MUTS2-2) in *Mitrastemon* may be linked to the unusual evolutionary trajectory of its plastome. The loss of both MUTS2 homologs is shared by another holoparasitic lineage with similar plastome features, the Balanophoraceae (Ceriotti et al., 2022; Schelkunov et al., 2021).

MUTS2-1 and MUTS2-2 originated from an ancient duplication predating the divergence of Viridiplantae (Sloan et al., 2025) and belong to the MutS mismatch repair family (Malik and Henikoff, 2000). However, accumulating evidence suggests that their primary function may not be directly related to canonical DNA repair. Studies in bacteria have implicated MutS2 homologs in recombination suppression (Pinto et al., 2005) and ribosomal stalling (Cerullo et al., 2022). Recent work in *Arabidopsis* mutants indicates that plastid-localized MUTS2 proteins participate in ribosome-associated quality control, particularly under conditions of high translational demand (Broz et al., 2025). Only limited evidence supports a role for these proteins in plastome homologous recombination (Broz et al., 2025). Taken together, the lack of MUTS2 expression in *Mitrastemon* may not directly compromise mismatch repair but could influence plastid genome stability indirectly through effects on plastid translational homeostasis, potentially contributing to the structural instability and AT enrichment observed in its plastome.

## FUNDING INFORMATION

Secretaría de Investigación, Internacionales y Posgrado, Universidad Nacional de Cuyo, Grant/Award #06/A092-T1 (to MVSP); Fondo para la Investigación Científica y Tecnológica, Grant/Award #PICT2020-01018 (to MVSP); National Natural Science Foundation of China (31811530297) (to R.Z.).

## DATA AVAILABILITY STATEMENT

Chloroplast sequence assembled and analyzed in this study is available in GenBank (PX991927), and electronic supplementary material is available online in Figshare (Roulet et al., 2026).

## AUTHOR CONTRIBUTION

MER and MVS-P conceived and designed the research. RY, CW, and RZ provided resources.

MER, LEG, and LG-S performed data analysis.

MER and MVS-P drafted and wrote the manuscript.

MER, LEG, LG-S, RZ, and MVS-P revised the manuscript.

## SUPPLEMENTARY DATA

**Figure S1. A**. DNA read depth of the plastid genomes calculated with Bowtie2 v.2.4.4 (parameters: --end-to-end --very-sensitive --no-contain --no-discordant --no-mixed). **B**. RNA read depth of the plastid genome and CDS genes of *Mitrastemon yamamotoi* #1 using RNA seq data from SRR28027597 (Carruthers et al., 2024) and calculated with Bowtie2 v.2.4.4 (parameters: --end-to-end --very-sensitive --no-discordant).

**Figure S2. A**. Linear representation of the circular plastid genomes of four *Mitrastemon* individuals. Genes are color-coded according to functional categories: large and small ribosomal subunits (LSU, SSU), ribosomal RNAs, tRNAs, and other genes. Genes containing cis-spliced introns are marked with asterisks. **B**. ProgressiveMauve whole-plastome alignment of *Mitrastemon* individuals. Locally Collinear Blocks (LCBs) are shown as colored regions connected across genomes, representing homologous segments shared among plastomes. Block orientation indicates structural collinearity. Gene maps are shown below each plastome, with functional categories indicated by color. Shorter *ycf* sequences reported in *M. yamamotoi* #3 remain uncertain due to the unavailability of the DNAseq data of this individual for read-based validation.

**Figure S3**. Maximum likelihood phylogenetic analysis of *Mitrastemon* based on a concatenated alignment of plastid (*clpP, rpl2, rpl16, rpl36, rps2, rps3, rps4, rps7, rps8, rps11, rps12, rps14, rps18*, and *rps19*) genes totaling 6,825 bp. Bootstrap support values >50% are shown above each branch. Scale bar corresponds to substitutions per site. Members of the family Ericales are shown in orange.

**Figure S4**. Root-to-tip substitution rates (dN and dS) across plastid genes in parasitic and autotrophic angiosperms. Scatterplots show synonymous (dS) and nonsynonymous (dN) substitution rates for each plastid gene separately, estimated from gene-specific phylogenies. Each point represents a terminal branch (tip) in the tree. Colors indicate different taxonomic groups: *Mitrastemon* (orange), photosynthetic Ericales (light green), and other autotrophic angiosperms (green).

**Figure S5**. Maximum likelihood phylogenies of individual plastid genes used for substitution rate analysis. Gene trees were reconstructed using IQ-TREE v.2.2.0 based on nucleotide alignments of 13 plastid genes retained across taxa. Branch labels indicate estimated substitution rates (dN or dS) as inferred by *codeml* under a branch model (PAML). These values represent root-to-tip evolutionary distances used in rate comparisons.

**Table S1**. List of species used in phylogenetic, substitution rate, and genome composition analyses.

**Table S2**. Repeats with >90% sequence identity detected in the *Mitrastemon* ptDNAs. Repeated sequences were identified using the get_repeats.sh script developed by Gandini et al. (2019).

**Table S3**. Features of plastid genes in angiosperm endoparasitic species.

**Table S4**. Summary of plastid substitution rate estimates (dN, dS, and dN/dS [ω]) for 13 protein-coding genes across endoparasitic and photosynthetic angiosperms.

**Table S5**. Plastid ribosomal protein subunits identified in the *Mitrastemon* transcriptome.

## Notes

https://doi.org/10.6084/m9.figshare.31802569

### Competing Interest Statement

The authors have declared no competing interest.

## LITERATURE CITED

Arias-Agudelo, L.M., González, F., Isaza, J.P., Alzate, J.F., Pabón-Mora, N., 2019. Plastome reduction and gene content in New World Pilostyles (Apodanthaceae) unveils high similarities to African and Australian congeners. Mol. Phylogenet. Evol. 135, 193–202. 10.1016/j.ympev.2019.03.014

Banerjee, A., Stefanović, S., 2023. A comparative study across the parasitic plants of Cuscuta subgenus Grammica (Convolvulaceae) reveals a possible loss of the plastid genome in its section Subulatae. Planta 257, 66. 10.1007/s00425-023-04099-y

Barbrook, A., Howe, C., Purton, S., 2006. Why are plastid genomes retained in non-photosynthetic organisms? Trends Plant Sci. 11, 101–8.

Bellot, S., Renner, S.S., 2016. The Plastomes of Two Species in the Endoparasite Genus Pilostyles (Apodanthaceae) Each Retain Just Five or Six Possibly Functional Genes. Genome Biol. Evol. 8, 189–201. 10.1093/gbe/evv251

Bromham, L., Cowman, P.F., Lanfear, R., 2013. Parasitic plants have increased rates of molecular evolution across all three genomes. BMC Evol. Biol. 13, 126. 10.1186/1471-2148-13-126

Broz, A.K., Kodrich, K., Prasad, K.V.S.K., Kuster, S.A., Lee, S., Forsythe, E.S., Sloan, D.B., 2025. Plant MutS2 proteins function in plastid ribosome quality control. 10.1101/2025.09.02.673837

Cai, L., 2023. Rethinking convergence in plant parasitism through the lens of molecular and population genetic processes. Am. J. Bot. 110, e16174. 10.1002/ajb2.16174

Cai, L., Arnold, B.J., Xi, Z., Khost, D.E., Patel, N., Hartmann, C.B., Manickam, S., Sasirat, S., Nikolov, L.A., Mathews, S., Sackton, T.B., Davis, C.C., 2021. Deeply Altered Genome Architecture in the Endoparasitic Flowering Plant Sapria himalayana Griff. (Rafflesiaceae). Curr. Biol. 31, 1002-1011.e9. 10.1016/j.cub.2020.12.045

Carruthers, T., Gonçalves, D.J.P., Li, P., Chanderbali, A.S., Dick, C.W., Fritsch, P.W., Larson, D.A., Soltis, D.E., Soltis, P.S., Weaver, W.N., Smith, S.A., 2024. Repeated shifts out of tropical climates preceded by whole genome duplication. New Phytol. 244, 2561–2575. 10.1111/nph.20200

Ceriotti, F., Roulet, M., Sanchez-Puerta, M.V., 2021. Plastomes in the holoparasitic family Balanophoraceae: Extremely high AT content, severe gene content reduction, and two independent genetic code changes. Mol. Phylogenet. Evol. 162, 107208.

Ceriotti, L.F., Gatica Soria, L.M., Guzman, S., Sato, H.A., Tovar Luque, E., Gonzalez, M.A., Sanchez-Puerta, M.V., 2025. The evolution of the plastid genomes in the holoparasitic Balanophoraceae. Proc. R. Soc. B Biol. Sci. 292, 20242011. 10.1098/rspb.2024.2011

Ceriotti, L.F., Gatica-Soria, L., Sanchez-Puerta, M.V., 2022. Cytonuclear coevolution in a holoparasitic plant with highly disparate organellar genomes. Plant Mol. Biol. 109, 673–688. 10.1007/s11103-022-01266-9

Cerullo, F., Filbeck, S., Patil, P.R., Hung, H.-C., Xu, H., Vornberger, J., Hofer, F.W., Schmitt, J., Kramer, G., Bukau, B., Hofmann, K., Pfeffer, S., Joazeiro, C.A.P., 2022. Bacterial ribosome collision sensing by a MutS DNA repair ATPase paralogue. Nature 603, 509–514. 10.1038/s41586-022-04487-6

Chen, X., Fang, D., Wu, C., Liu, B., Liu, Y., Sahu, S.K., Song, B., Yang, S., Yang, T., Wei, J., Wang, X., Zhang, W., Xu, Q., Wang, H., Yuan, L., Liao, X., Chen, L., Chen, Z., Yuan, F., Chang, Y., Lu, L., Yang, H., Wang, J., Xu, X., Liu, X., Wicke, S., Liu, H., 2020. Comparative Plastome Analysis of Root- and Stem-Feeding Parasites of Santalales Untangle the Footprints of Feeding Mode and Lifestyle Transitions. Genome Biol. Evol. 12, 3663–3676. 10.1093/gbe/evz271

Doyle, J., 1991. DNA Protocols for Plants, in: Hewitt, G.M., Johnston, A.W.B., Young, J.P.W. (Eds.), Molecular Techniques in Taxonomy. Springer, Berlin, Heidelberg, pp. 283–293. 10.1007/978-3-642-83962-7_18

Gandini, C.L., Garcia, L., Abbona, C.C., Sanchez-Puerta, M.V., 2019. The complete organelle genomes of Physochlaina orientalis: insights into short sequence repeats across seed plant mitochondrial genomes. Mol. Phylogenet. Evol. 137, 274–284.

Gordon, D., Green, P., 2013. Consed: a graphical editor for next-generation sequencing. Bioinformatics 29, 2936–2937. 10.1093/bioinformatics/btt515

Grabherr, M.G., Haas, B.J., Yassour, M., Levin, J.Z., Thompson, D.A., Amit, I., Adiconis, X., Fan, L., Raychowdhury, R., Zeng, Q., Chen, Z., Mauceli, E., Hacohen, N., Gnirke, A., Rhind, N., di Palma, F., Birren, B.W., Nusbaum, C., Lindblad-Toh, K., Friedman, N., Regev, A., 2011. Full-length transcriptome assembly from RNA-Seq data without a reference genome. Nat. Biotechnol. 29, 644–652. 10.1038/nbt.1883

Graham, S.W., Lam, V.K.Y., Merckx, V.S.F.T., 2017. Plastomes on the edge: the evolutionary breakdown of mycoheterotroph plastid genomes. New Phytol. 214, 48– 55. 10.1111/nph.14398

Gualberto, J.M., Newton, K.J., 2017. Plant Mitochondrial Genomes: Dynamics and Mechanisms of Mutation. Annu. Rev. Plant Biol. 68, 225–252. 10.1146/annurev-arplant-043015-112232

Jin, J.-J., Yu, W.-B., Yang, J.-B., Song, Y., dePamphilis, C.W., Yi, T.-S., Li, D.-Z., 2020. GetOrganelle: a fast and versatile toolkit for accurate de novo assembly of organelle genomes. Genome Biol. 21, 241. 10.1186/s13059-020-02154-5

Katoh, K., Standley, D.M., 2013. MAFFT Multiple Sequence Alignment Software Version 7: Improvements in Performance and Usability. Mol. Biol. Evol. 30, 772– 780. 10.1093/molbev/mst010

Kearse, M., Moir, R., Wilson, A., Stones-Havas, S., Cheung, M., Sturrock, S., Buxton, S., Cooper, A., Markowitz, S., Duran, C., Thierer, T., Ashton, B., Meintjes, P., Drummond, A., 2012. Geneious Basic: An integrated and extendable desktop software platform for the organization and analysis of sequence data. Bioinformatics 28, 1647–1649. 10.1093/bioinformatics/bts199

Langmead, B., Salzberg, S.L., 2012. Fast gapped-read alignment with Bowtie 2. Nat. Methods 9, 357–359. 10.1038/nmeth.1923

Liu, X., Liao, X., Chen, D., Zheng, Y., Yu, X., Xu, X., Liu, Z., Lan, S., 2020. The complete chloroplast genome sequence of Monotropa uniflora (Ericaceae). Mitochondrial DNA Part B Resour. 5, 3168–3169. 10.1080/23802359.2020.1806754

Malik, H.S., Henikoff, S., 2000. Dual recognition–incision enzymes might be involved in mismatch repair and meiosis. Trends Biochem. Sci. 25, 414–418. 10.1016/S0968-0004(00)01623-6

Maréchal, A., Brisson, N., 2010. Recombination and the maintenance of plant organelle genome stability. New Phytol. 186, 299–317. 10.1111/j.1469-8137.2010.03195.x

Minh, B.Q., Schmidt, H.A., Chernomor, O., Schrempf, D., Woodhams, M.D., von Haeseler, A., Lanfear, R., 2020. IQ-TREE 2: New Models and Efficient Methods for Phylogenetic Inference in the Genomic Era. Mol. Biol. Evol. 37, 1530–1534. 10.1093/molbev/msaa015

Molina, J., Hazzouri, K.M., Nickrent, D.L., Geisler, M., Meyer, R.S., Pentony, M.M., Flowers, J.M., Pelser, P., Barcelona, J., Inovejas, S.A., Uy, I., Yuan, W., Wilkins, O., Michel, C.I., LockLear, S., Concepcion, G.P., Purugganan, M., 2014. Possible loss of the chloroplast genome in the parasitic flowering plant Rafflesia lagascae (Rafflesiaceae). Mol. Biol. Evol. 31, 793–803.

Naumann, J., Der, J.P., Wafula, E., Jones, S., Wagner, S., Honaas, L., Ralph, P., Bolin, J., Maass, E., Neinhuis, C., Wanke, S., dePamphilis, C.W., 2016. Detecting and characterizing the highly divergent plastid genome of the nonphotosynthetic parasitic plant Hydnora visseri (Hydnoraceae). Genome Biol Evol 8, 345–363.

Nickrent, D.L., 2020. Parasitic angiosperms: how often and how many? Taxon 69, 5– 27.

Nickrent, D.L., Starr, E.M., 1994. High rates of nucleotide substitution in nuclear small-subunit (18S) rDNA from holoparasitic flowering plants. J. Mol. Evol. 39, 62– 70. 10.1007/BF00178250

Odahara, M., Masuda, Y., Sato, M., Wakazaki, M., Harada, C., Toyooka, K., Sekine, Y., 2015. RECG Maintains Plastid and Mitochondrial Genome Stability by Suppressing Extensive Recombination between Short Dispersed Repeats. PLOS Genet. 11, e1005080. 10.1371/journal.pgen.1005080

Pinto, A.V., Mathieu, A., Marsin, S., Veaute, X., Ielpi, L., Labigne, A., Radicella, J.P., 2005. Suppression of Homologous and Homeologous Recombination by the Bacterial MutS2 Protein. Mol. Cell 17, 113–120. 10.1016/j.molcel.2004.11.035

Ramírez-Ramírez, J.A., González, F., Elejalde-Baena, E., Alzate, J.F., Pabón-Mora, N., 2025. Secrets within stems: The cryptic Apodanthes caseariae (Apodanthaceae), a rare neotropical holoendoparasite. PLANTS PEOPLE PLANET n/a. 10.1002/ppp3.70102

Roquet, C., Coissac, É., Cruaud, C., Boleda, M., Boyer, F., Alberti, A., Gielly, L., Taberlet, P., Thuiller, W., Van Es, J., Lavergne, S., 2016. Understanding the evolution of holoparasitic plants: the complete plastid genome of the holoparasite Cytinus hypocistis (Cytinaceae). Ann. Bot. 118, 885–896. 10.1093/aob/mcw135

Roulet, M., Gatica-Soria, L., Garcia, L., Yu, R., Wang, C., Zhou, R., Sanchez-Puerta, M., 2026. Supplementary material from: Plastome convergence across parasitic lineages: genome reduction, extreme AT bias, and functional persistence in the endoparasitic Mitrastemonaceae. 10.6084/m9.figshare.31802569

Roulet, M.E., Garcia, L.E., Yu, R., Wang, C., Zhou, R., Sanchez-Puerta, M.V., 2025. Circle-Mediated HGT shapes the multichromosomal mitochondrial genome of the endoparasite Mitrastemon yamamotoi. 10.1101/2025.06.25.661533

Sanchez-Puerta, M., Ceriotti, L., Gatica-Soria, L., Roulet, M., Garcia, L.E., Sato, H., 2023. Beyond parasitic convergence: unraveling the evolution of the organellar genomes in holoparasites. Ann. Bot. (in press) doi: 10.1093/aob/mcad108. https://doi.org/10.1093/aob/mcad108

Scarpin, M.R., Busche, M., Martinez, R.E., Harper, L.C., Reiser, L., Szakonyi, D., Merchante, C., Lan, T., Xiong, W., Mo, B., Tang, G., Chen, X., Bailey-Serres, J., Browning, K.S., Brunkard, J.O., 2023. An updated nomenclature for plant ribosomal protein genes. Plant Cell 35, 640–643. 10.1093/plcell/koac333

Schelkunov, M., Nuraliev, M., Logacheva, M., 2021. Genomic comparison of non-photosynthetic plants from the family Balanophoraceae with their photosynthetic relatives. PeerJ. 9, e12106.

Sloan, D.B., Broz, A.K., Kuster, S.A., Muthye, V., Peñafiel-Ayala, A., Marron, J.R., Lavrov, D.V., Brieba, L.G., 2025. Expansion of the MutS gene family in plants. Plant Cell 37, koae277. 10.1093/plcell/koae277

Su, H.-J., Barkman, T.J., Hao, W., Jones, S., Naumann, J., Skippington, E., Wafula, E., Hu, J.-M., Palmer, J.D., dePamphilis, C.W., 2019. Novel genetic code and record-setting AT-richness in the highlly reduced plastid genome of the holoparasitic plant Balanophora. Proc. Natl. Acad. Sci. 116, 934–943.

Teixeira-Costa, L., Suetsugu, K., 2023. Neglected plant parasites: Mitrastemonaceae. PLANTS PEOPLE PLANET 5, 5–13. 10.1002/ppp3.10322

The Angiosperm Phylogeny Group, Chase, M.W., Christenhusz, M.J.M., Fay, M.F., Byng, J.W., Judd, W.S., Soltis, D.E., Mabberley, D.J., Sennikov, A.N., Soltis, P.S., Stevens, P.F., 2016. An update of the Angiosperm Phylogeny Group classification for the orders and families of flowering plants: APG IV. Bot. J. Linn. Soc. 181, 1–20. 10.1111/boj.12385

Tillich, M., Lehwark, P., Pellizzer, T., Ulbricht-Jones, E.S., Fischer, A., Bock, R., Greiner, S., 2017. GeSeq – versatile and accurate annotation of organelle genomes. Nucleic Acids Res. 45, W6–W11. 10.1093/nar/gkx391

Wick, R.R., Schultz, M.B., Zobel, J., Holt, K.E., 2015. Bandage: interactive visualization of de novo genome assemblies. Bioinformatics 31, 3350–3352. 10.1093/bioinformatics/btv383

Wicke, S., Muller, K.F., de Pamphilis, C.W., Quandt, D., Wickett, N., Zhang, Y., Renner, S.S., Schneeweiss, G.M., 2013. Mechanisms of functional and physical genome reduction in photosynthetic and nonphotosynthetic parasitic plants of the broomrape family. Plant Cell 25, 3711–3725.

Wicke, S., Naumann, J., 2018. Chapter Eleven - Molecular Evolution of Plastid Genomes in Parasitic Flowering Plants, in: Chaw, S.-M., Jansen, R.K. (Eds.), Advances in Botanical Research. Academic Press, pp. 315–347. 10.1016/bs.abr.2017.11.014

Wu, Z., Waneka, G., Broz, A.K., King, C.R., Sloan, D.B., 2020. MSH1 is required for maintenance of the low mutation rates in plant mitochondrial and plastid genomes. Proc. Natl. Acad. Sci. 117, 16448–16455. 10.1073/pnas.2001998117

Yang, Z., 2007. PAML 4: Phylogenetic Analysis by Maximum Likelihood. Mol. Biol. Evol. 24, 1586–1591. 10.1093/molbev/msm088

Yu, R., Zhi, X., Ceriotti, L.F., Skippington, E., Rice, D.W., Su, H.-J., Barkman, T.J., Sun, C., Liu, Y., Fang, D., Chen, X., dePamphilis, C.W., Mower, J.P., Sanchez-Puerta, M.V., Palmer, J.D., Zhou, R., 2025. A record-setting mitogenome in the holoparasitic plant Balanophora yakushimensis accompanied by exceptional loss of organellar DNA repair and recombination genes. BMC Biol. 23, 344. 10.1186/s12915-025-02449-8

Zhang, J., Ruhlman, T.A., Sabir, J.S.M., Blazier, J.C., Weng, M.-L., Park, S., Jansen, R.K., 2016. Coevolution between Nuclear-Encoded DNA Replication, Recombination, and Repair Genes and Plastid Genome Complexity. Genome Biol. Evol. 8, 622–634. 10.1093/gbe/evw033

