## Supplementary Figures for "Plastome convergence across heterotrophic plant lineages: genome reduction, extreme AT bias, high substitution rates, and functional persistence in the endoparasitic Mitrastemonaceae"

**Figure S1. A.** DNA read depth of the plastid genomes calculated with Bowtie2 v.2.4.4 (parameters: --end-to-end --very-sensitive --no-contain --no-discordant --no-mixed). **B.** RNA read depth of the plastid genome and CDS genes of *Mitrastemon yamamotoi* #1 using RNA seq data from SRR28027597 (Carruthers et al., 2024) and calculated with Bowtie2 v.2.4.4 (parameters: --end-to-end --very-sensitive --no-discordant).

**A.**

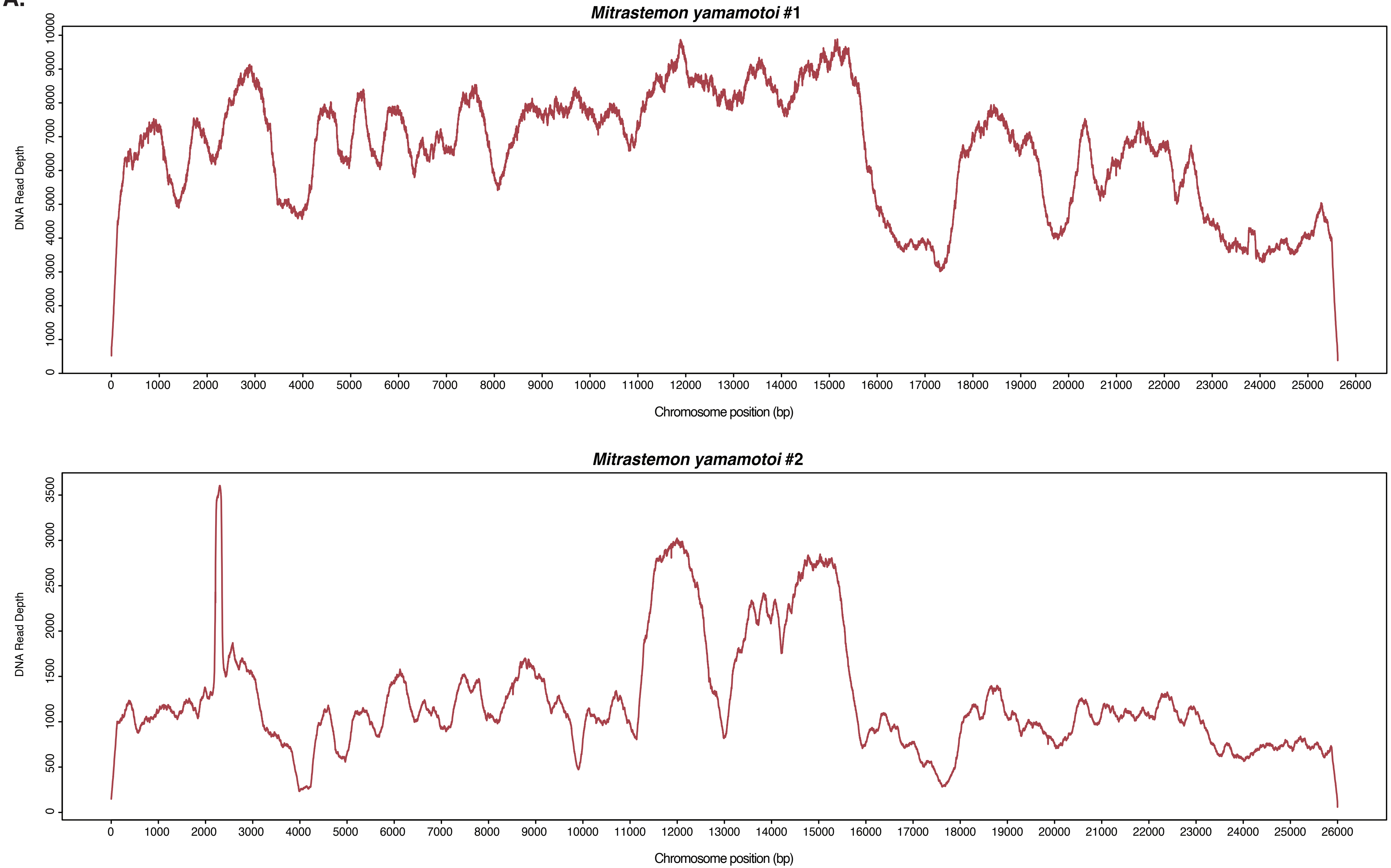

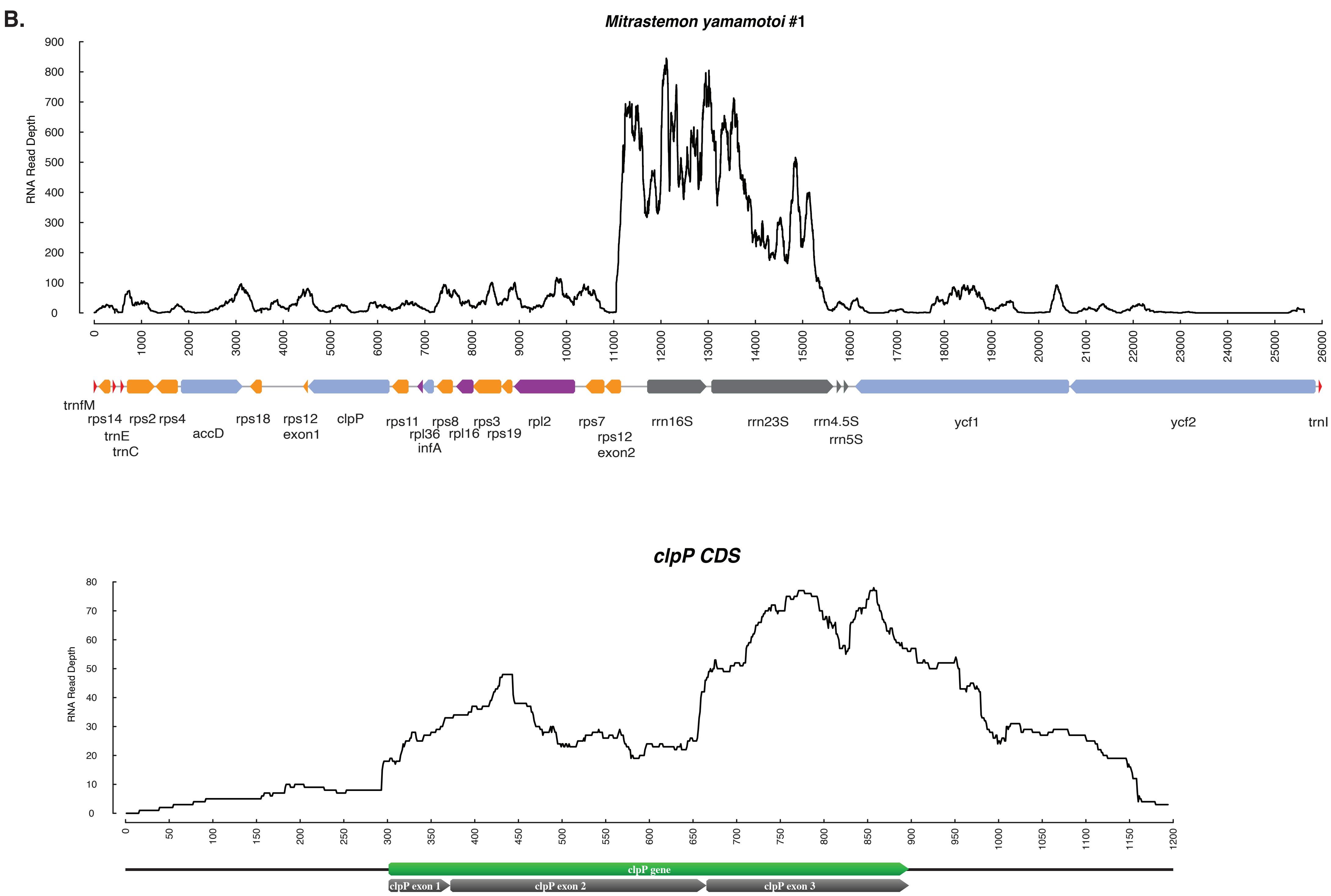
B.
Mitrastemon yamamotoi #1
RNA Read Depth
0
100
200
300
400
500
600
700
800
900
0
1000
2000
3000
4000
5000
6000
7000
8000
9000
10000
11000
12000
13000
14000
15000
16000
17000
18000
19000
20000
21000
22000
23000
24000
25000
26000
trnfM
rps14
trnE
trnC
rps2
rps4
accD
rps18
rps12 exon1
clpP
rps11
rpl36
infA
rps8
rpl16
rps3
rps19
rpl2
rps7
rps12 exon2
rrn16S
rrn23S
rrn4.5S
rrn5S
ycf1
ycf2
trnI
clpP CDS
RNA Read Depth
0
10
20
30
40
50
60
70
80
0
50
100
150
200
250
300
350
400
450
500
550
600
650
700
750
800
850
900
950
1000
1050
1100
1150
1200
clpP gene
clpP exon 1
clpP exon 2
clpP exon 3

#### *rpl2 CDS*

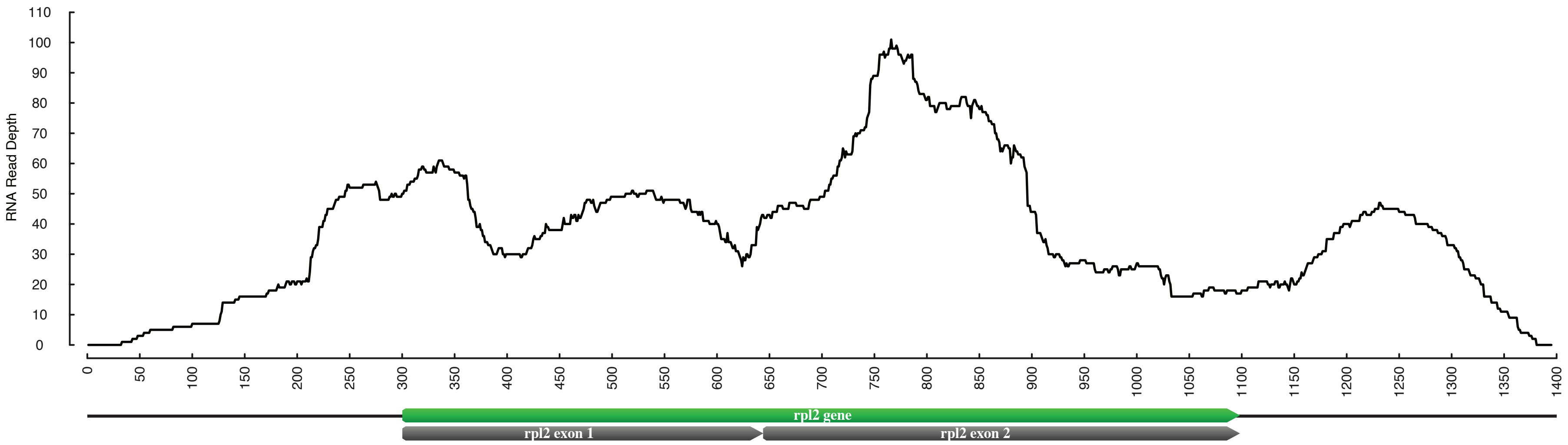


#### *rps12 CDS*

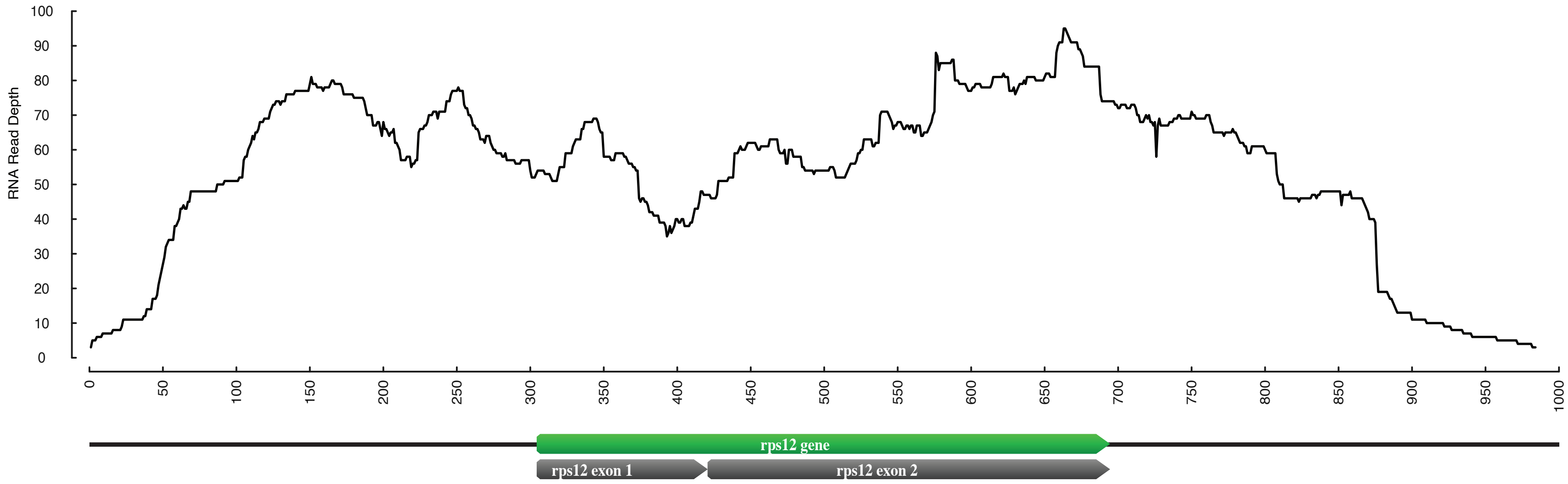

**Figure S2**. **A.** Linear representation of the circular plastid genomes of four *Mitrastemon* individuals. Genes are color-coded according to functional categories: large and small ribosomal subunits (LSU, SSU), ribosomal RNAs, tRNAs, and other genes. Genes containing cis-spliced introns are marked with asterisks. **B.** ProgressiveMauve whole-plastome alignment of *Mitrastemon* individuals. Locally Collinear Blocks (LCBs) are shown as colored regions connected across genomes, representing homologous segments shared among plastomes. Block orientation indicates structural collinearity. Gene maps are shown below each plastome, with functional categories indicated by color. Shorter *ycf* sequences reported in *M. yamamotoi* #3 remain uncertain due to the unavailability of the DNAseq data of this individual for read-based validation.

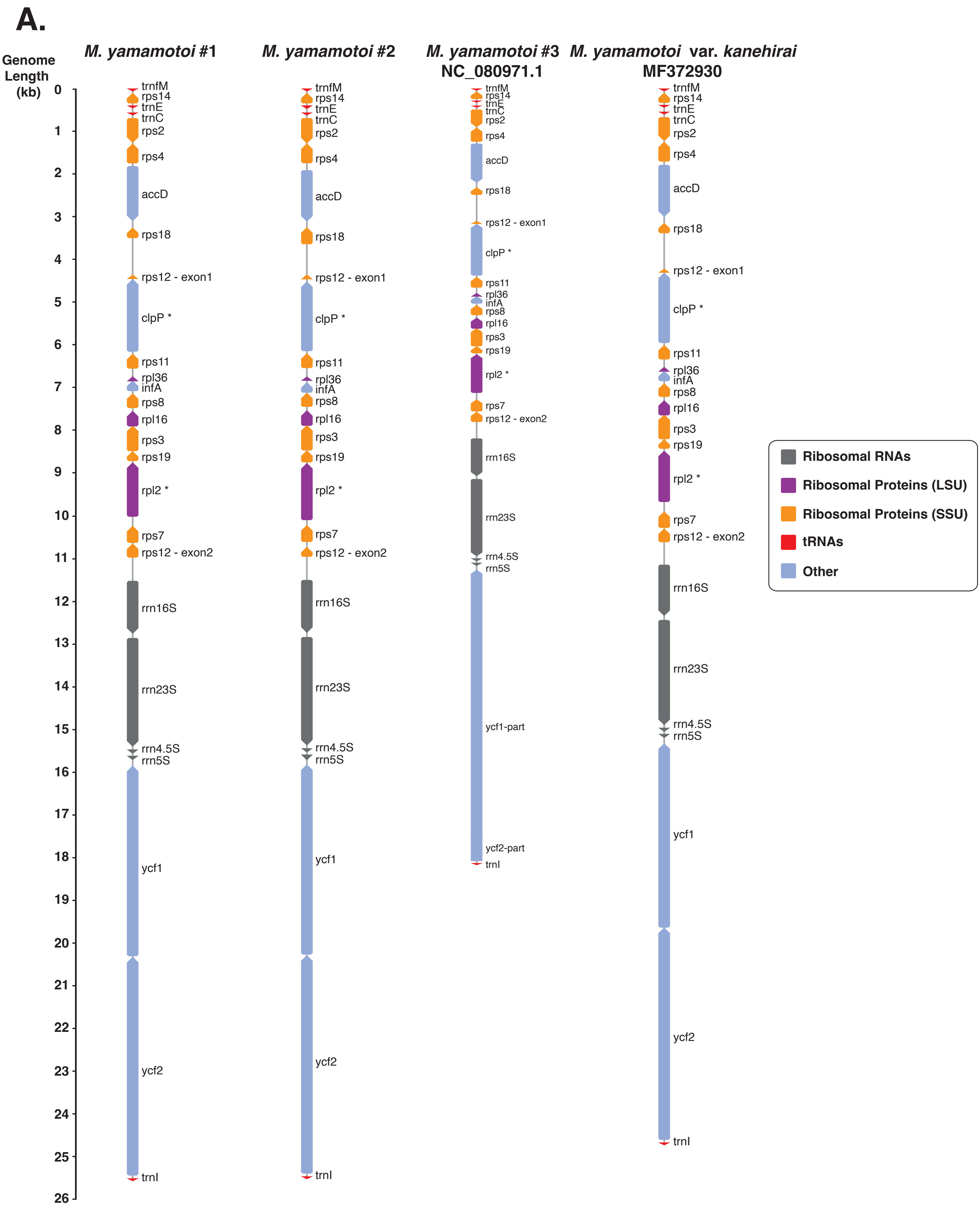

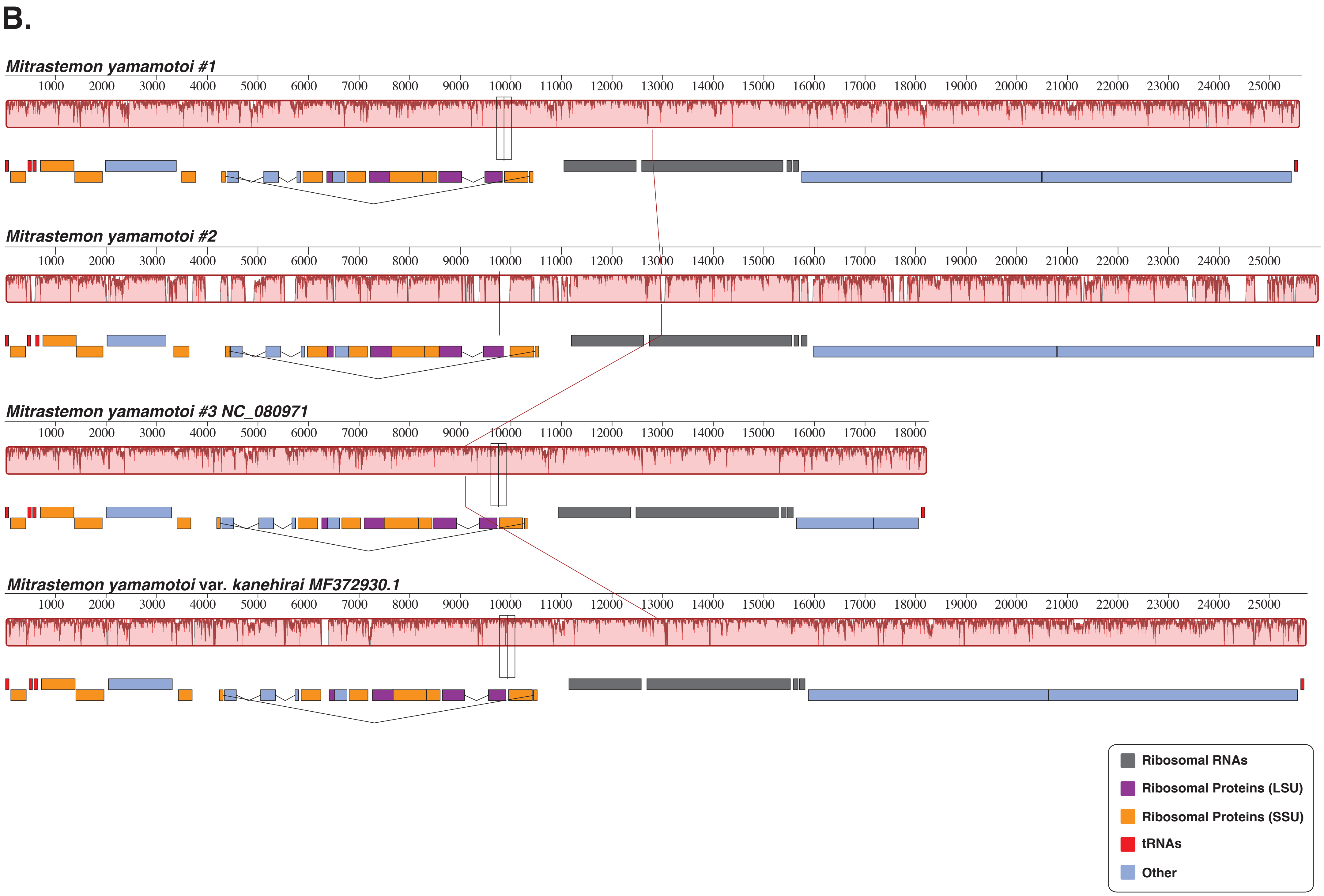

B.
Mitrastemon yamamotoi #1
1000 2000 3000 4000 5000 6000 7000 8000 9000 10000 11000 12000 13000 14000 15000 16000 17000 18000 19000 20000 21000 22000 23000 24000 25000
Mitrastemon yamamotoi #2
1000 2000 3000 4000 5000 6000 7000 8000 9000 10000 11000 12000 13000 14000 15000 16000 17000 18000 19000 20000 21000 22000 23000 24000 25000
Mitrastemon yamamotoi #3 NC_080971
1000 2000 3000 4000 5000 6000 7000 8000 9000 10000 11000 12000 13000 14000 15000 16000 17000 18000
Mitrastemon yamamotoi var. kanehirai MF372930.1
1000 2000 3000 4000 5000 6000 7000 8000 9000 10000 11000 12000 13000 14000 15000 16000 17000 18000 19000 20000 21000 22000 23000 24000 25000
Ribosomal RNAs
Ribosomal Proteins (LSU)
Ribosomal Proteins (SSU)
tRNAs
Other

**Figure S3**. **Maximum likelihood phylogenetic analysis of *Mitrastemon* based on a concatenated alignment of plastid (*clpP, rpl2, rpl16, rpl36, rps2, rps3, rps4, rps7, rps8, rps11, rps12, rps14, rps18,* and *rps19*) genes totaling 6,825 bp.** Bootstrap support values >50% are shown above each branch. Scale bar corresponds to substitutions per site. Members of the family Ericales are shown in orange.

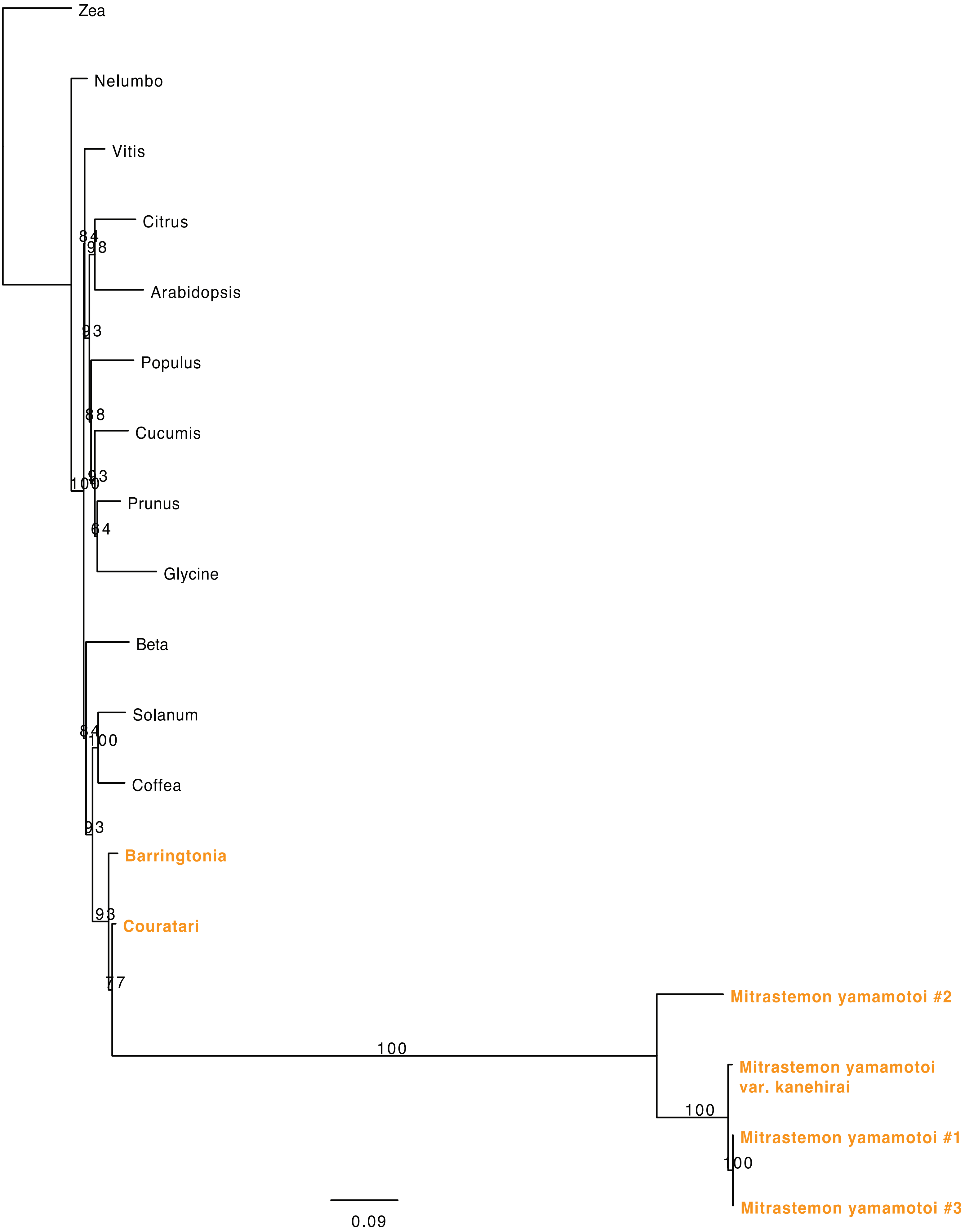

**Figure S4. Root-to-tip substitution rates (dN and dS) across plastid genes in parasitic and autotrophic angiosperms.** Scatterplots show synonymous (dS) and nonsynonymous (dN) substitution rates for each plastid gene separately, estimated from gene-specific phylogenies. Each point represents a terminal branch (tip) in the tree. Colors indicate different taxonomic groups: *Mitrastemon* (orange), photosynthetic Ericales (lightgreen), and other autotrophic angiosperms (green).

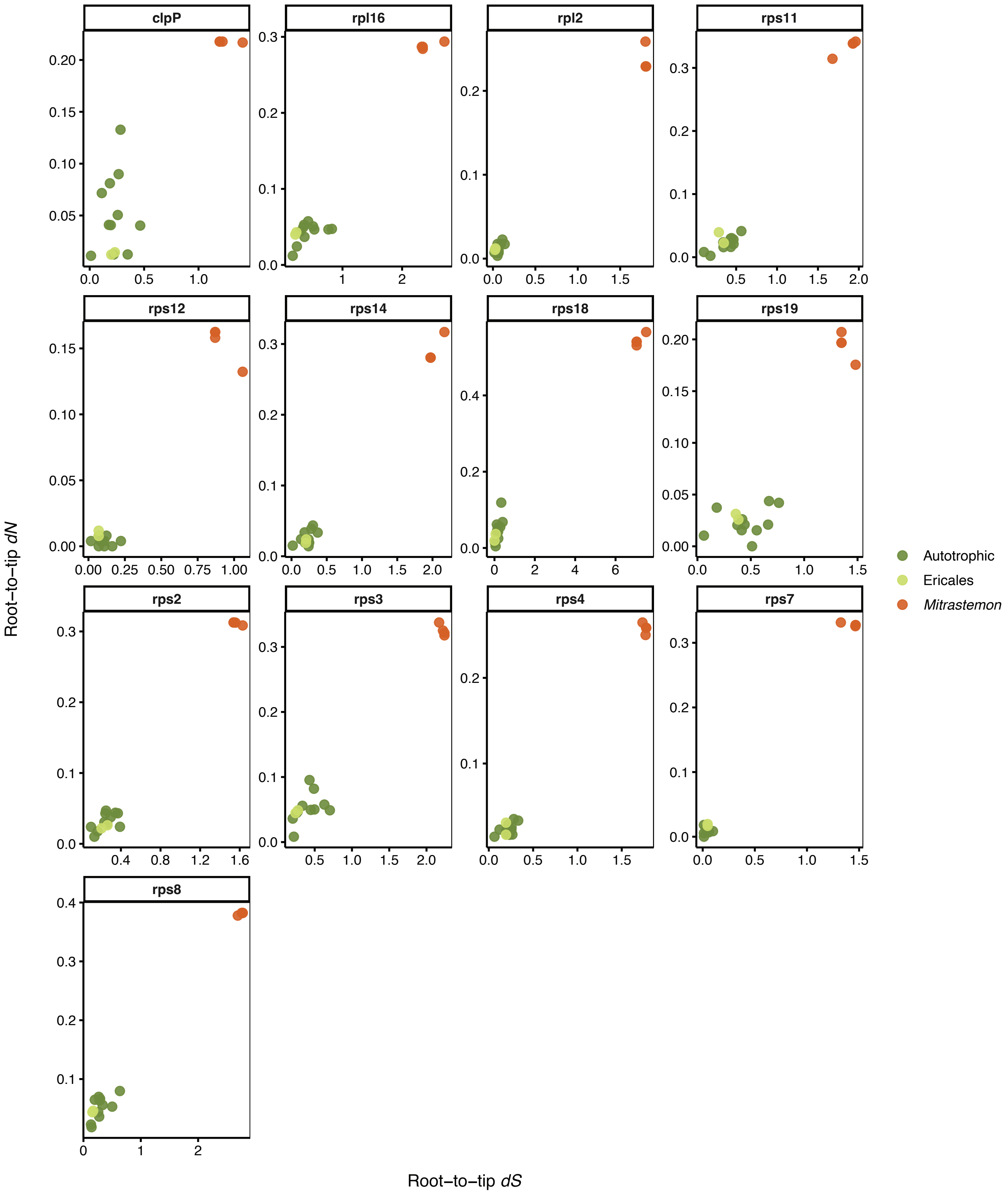

**Figure S5. Maximum likelihood phylogenies of individual plastid genes used for substitution rate analysis.** Gene trees were reconstructed using IQ-TREE v.2.2.0 based on nucleotide alignments of 13 plastid genes retained across taxa. Branch labels indicate estimated substitution rates (dN or dS) as inferred by codeml under a branch model (PAML). These values represent root-to-tip evolutionary distances used in rate comparisons.

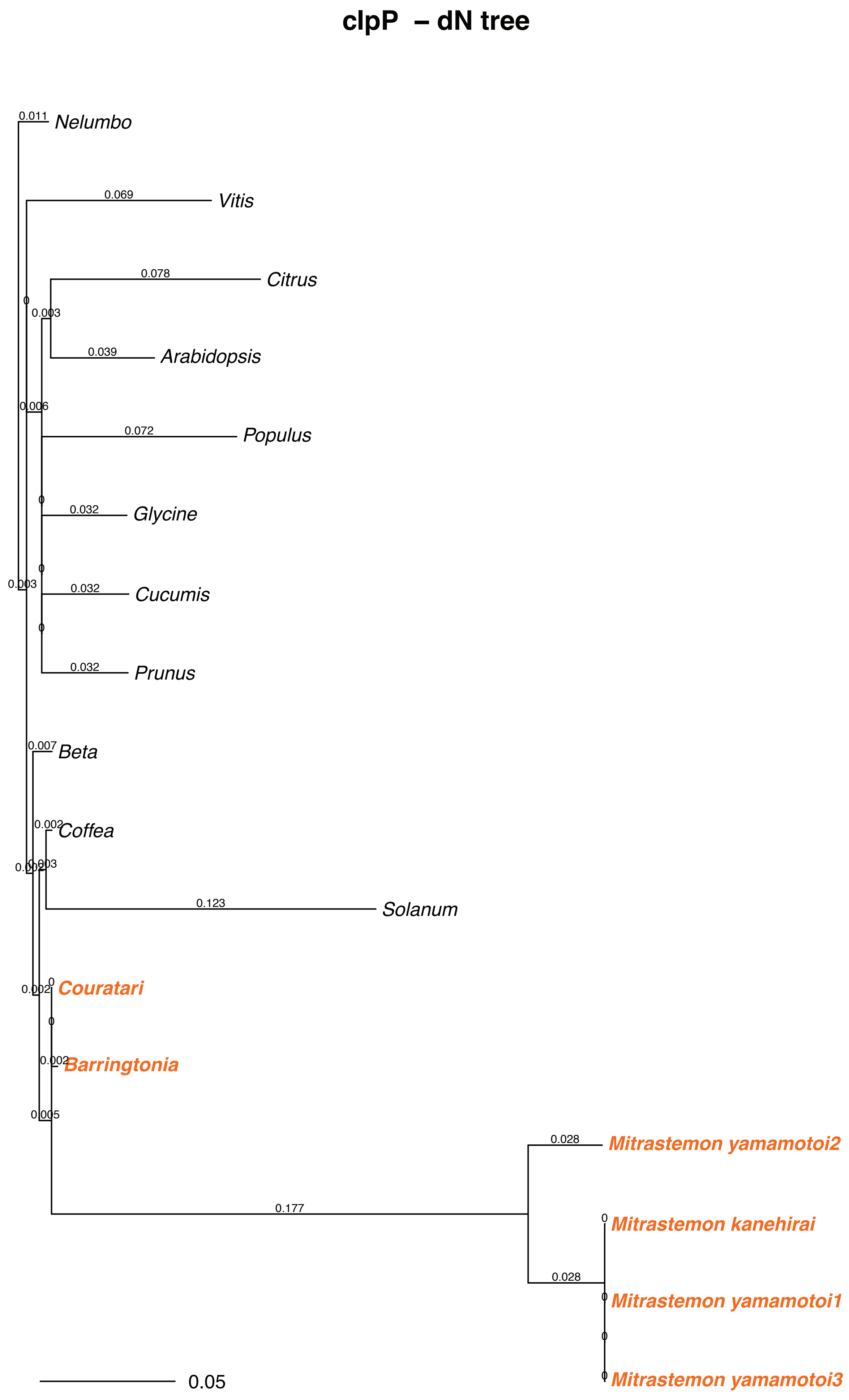

### clpP – dS tree

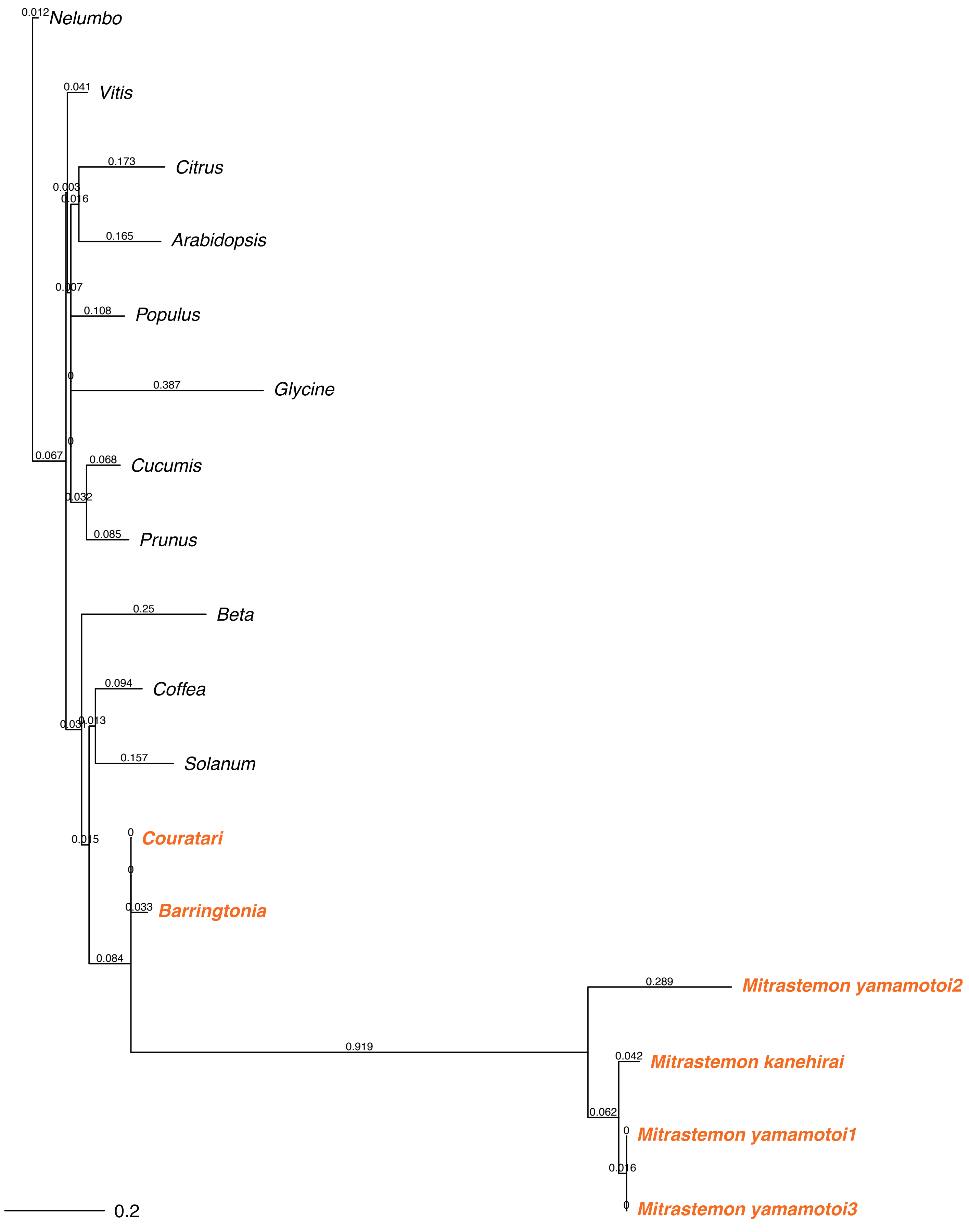

0.012
Nelumbo
0.041
Vitis
0.173
Citrus
0.003
0.016
0.165
Arabidopsis
0.007
0.108
Populus
0
0.387
Glycine
0
0.067
0.068
Cucumis
0.032
0.085
Prunus
0.25
Beta
0.094
Coffea
0.031
0.013
0.157
Solanum
0
0.015
Couratari
0
0.033
Barringtonia
0.084
0.289
Mitrastemon yamamotoi2
0.919
0.042
Mitrastemon kanehirai
0.062
0
Mitrastemon yamamotoi1
0.016
0
Mitrastemon yamamotoi3
0.2

**rpl2 – dN tree**

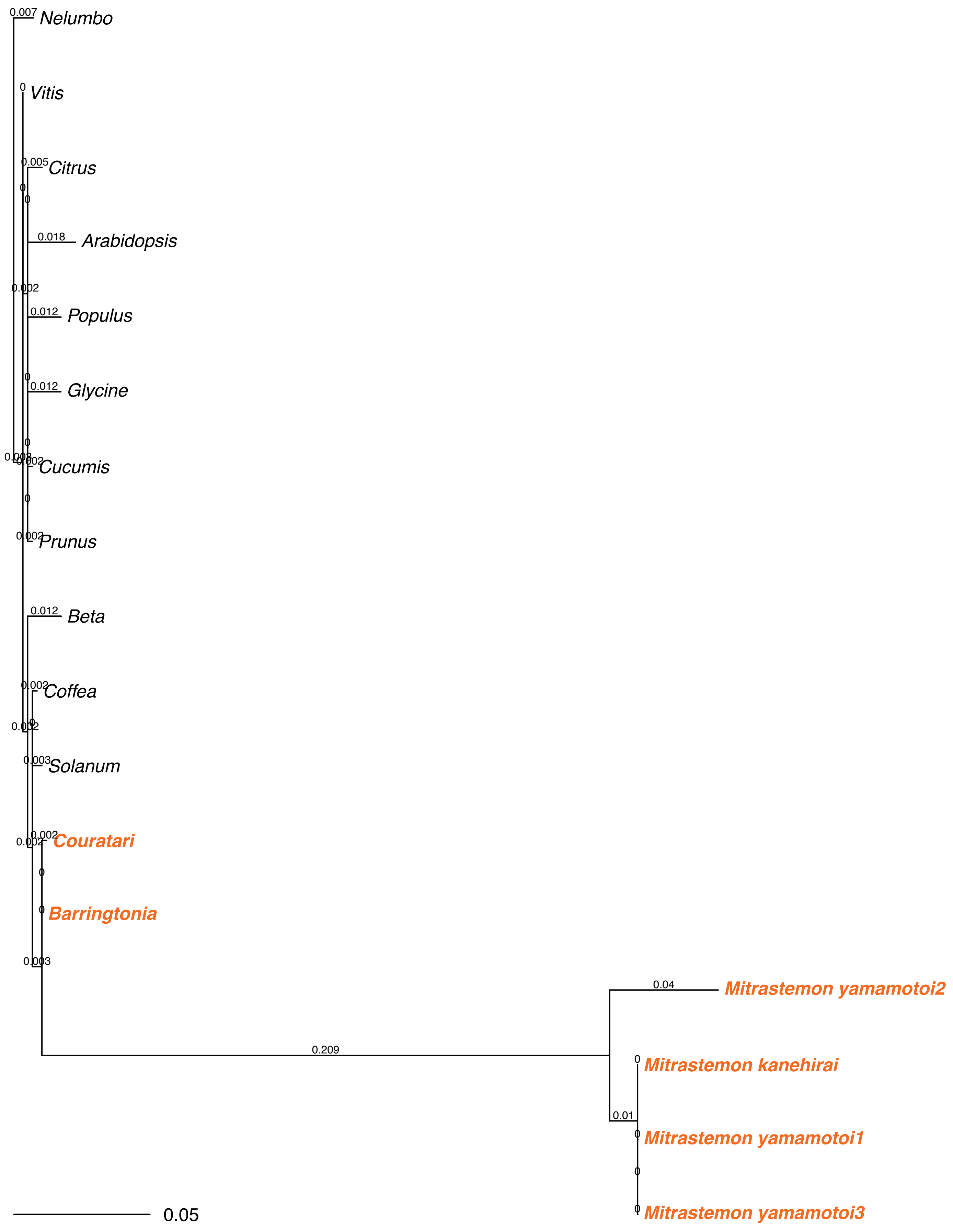

0.007 Nelumbo
0 Vitis
0.005 Citrus
0.018 Arabidopsis
0.012 Populus
0.012 Glycine
Cucumis
Prunus
0.012 Beta
Coffea
Solanum
Couratari
Barringtonia
0.209
0.04 Mitrastemon yamamotoi2
0 Mitrastemon kanehirai
0.01
Mitrastemon yamamotoi1
Mitrastemon yamamotoi3
0.05

### rpl2 – dS tree

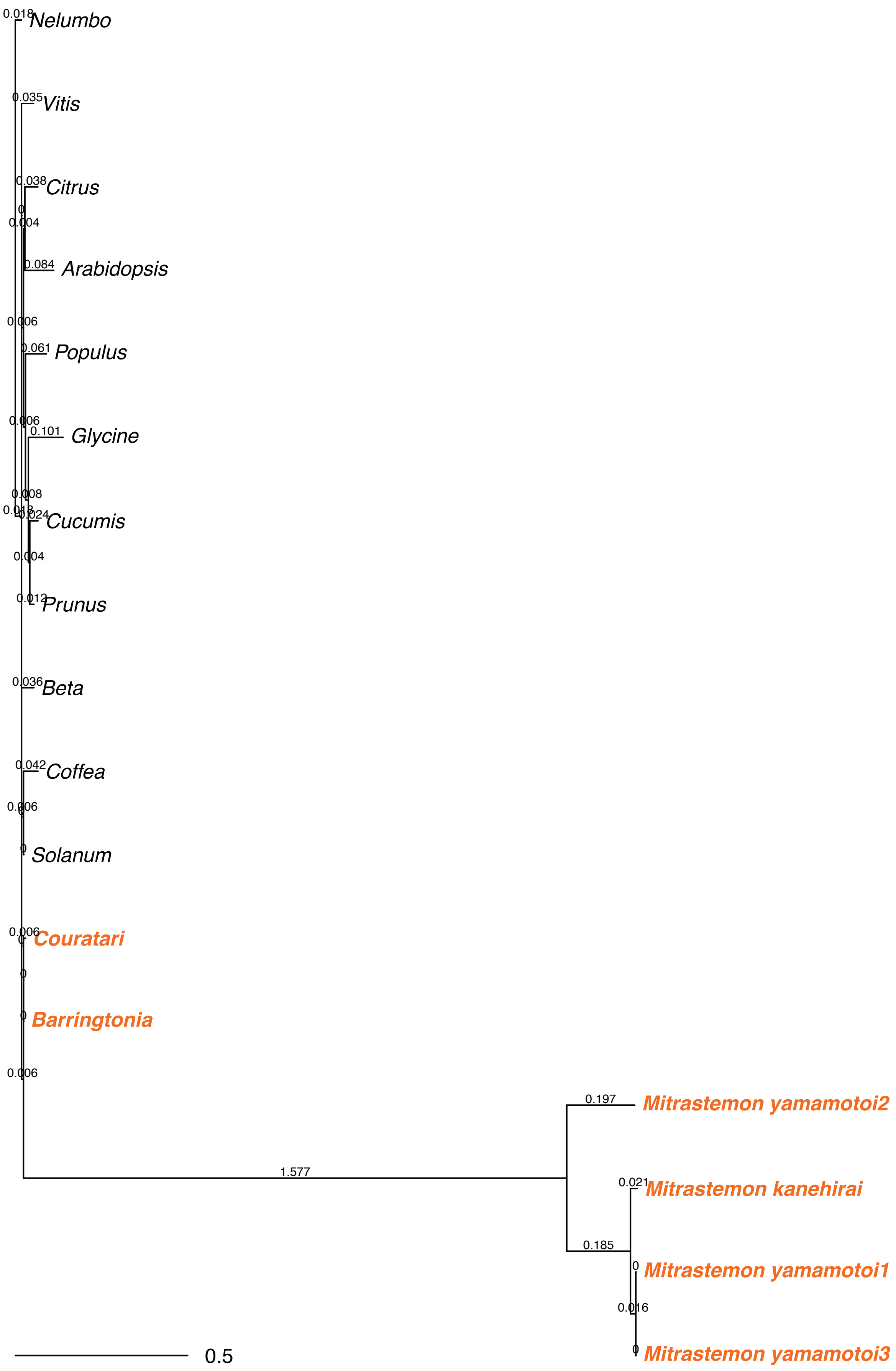
Nelumbo
Vitis
Citrus
Arabidopsis
Populus
Glycine
Cucumis
Prunus
Beta
Coffea
Solanum
Couratari
Barringtonia
Mitrastemon yamamotoi2
Mitrastemon kanehirai
Mitrastemon yamamotoi1
Mitrastemon yamamotoi3
0.018
0.035
0.038
0.004
0.084
0.006
0.061
0.101
0.008
0.024
0.004
0.197
1.577
0.021
0.185
0.016
0.5

#### rpl16 – dN tree

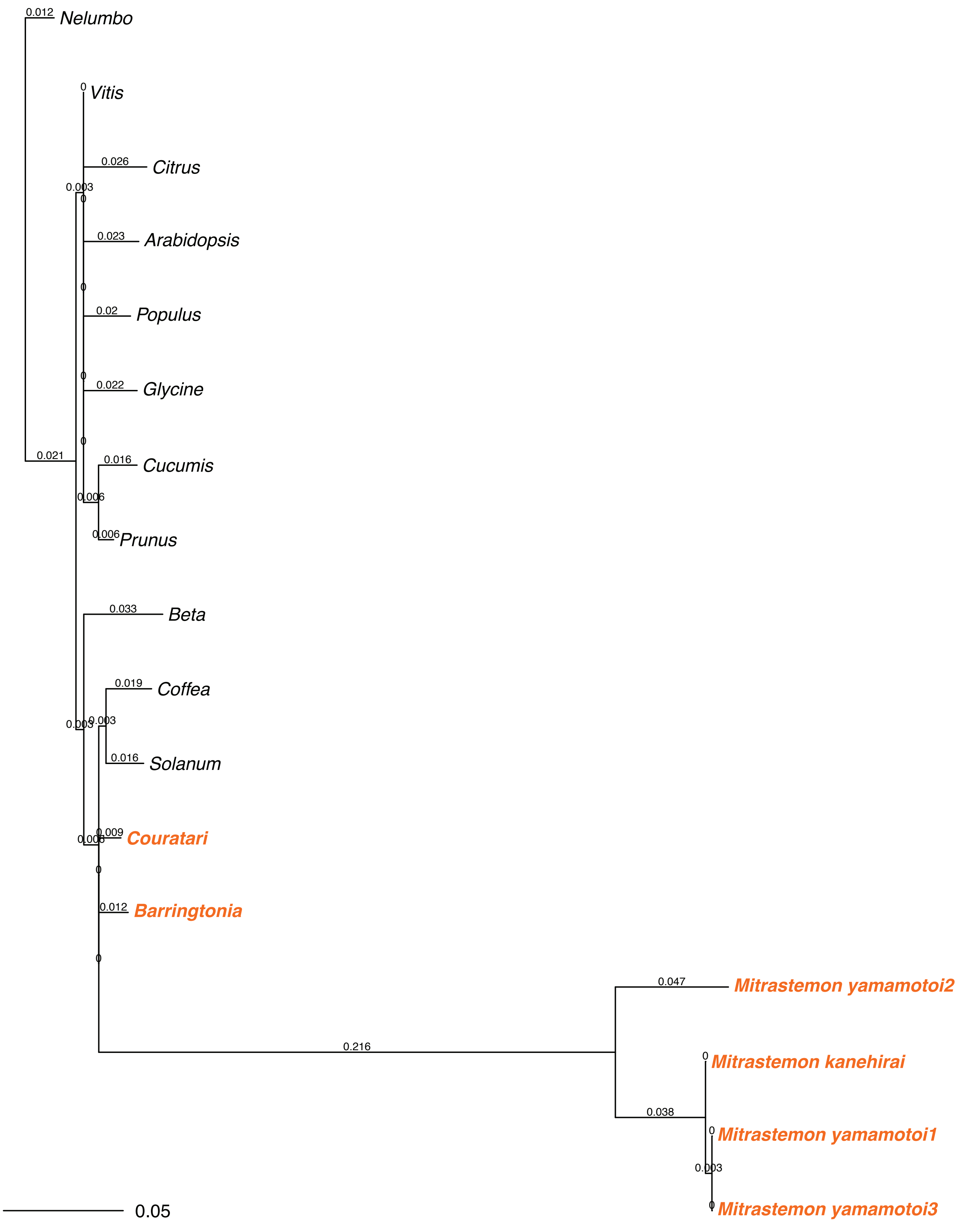

### rpl16 – dS tree

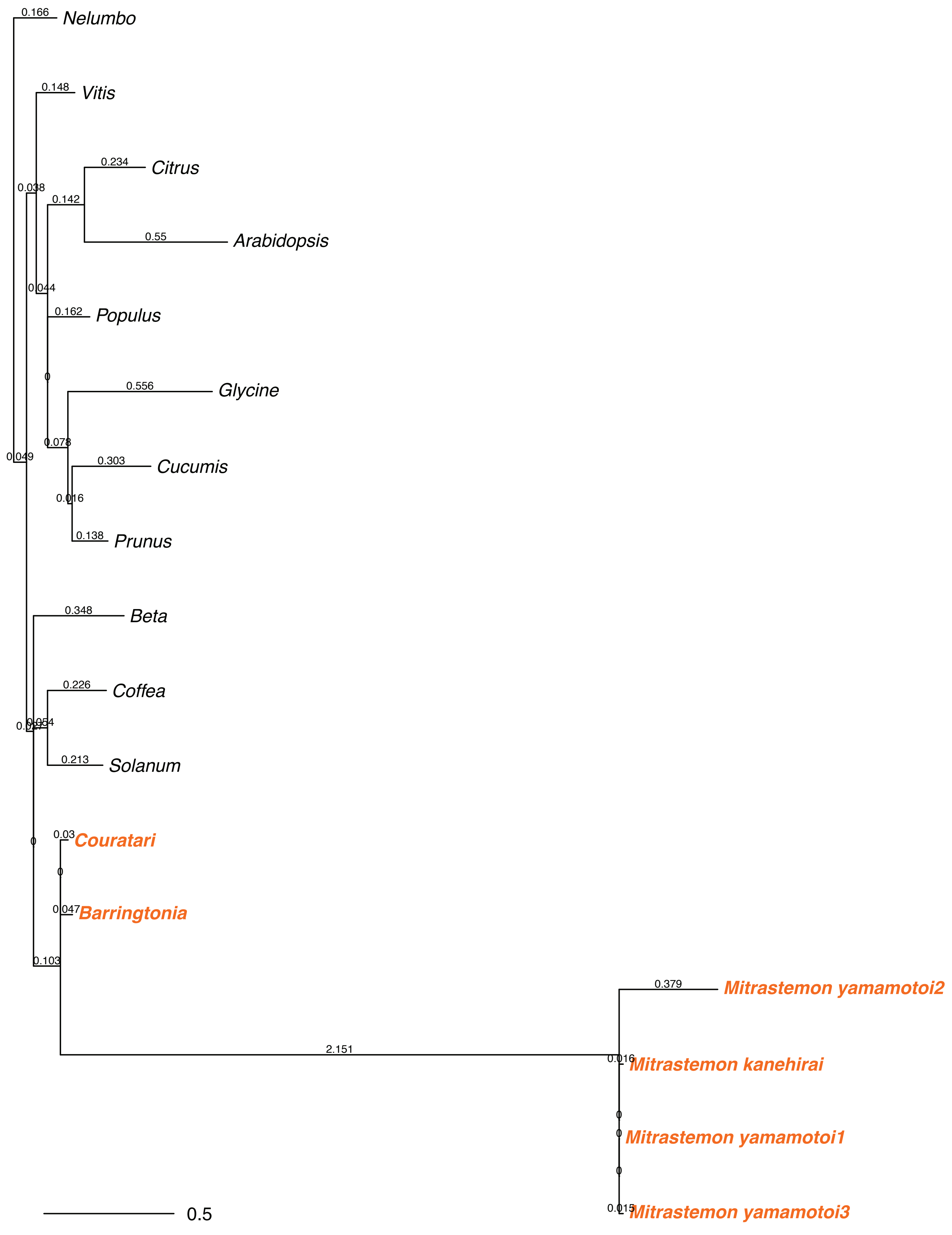

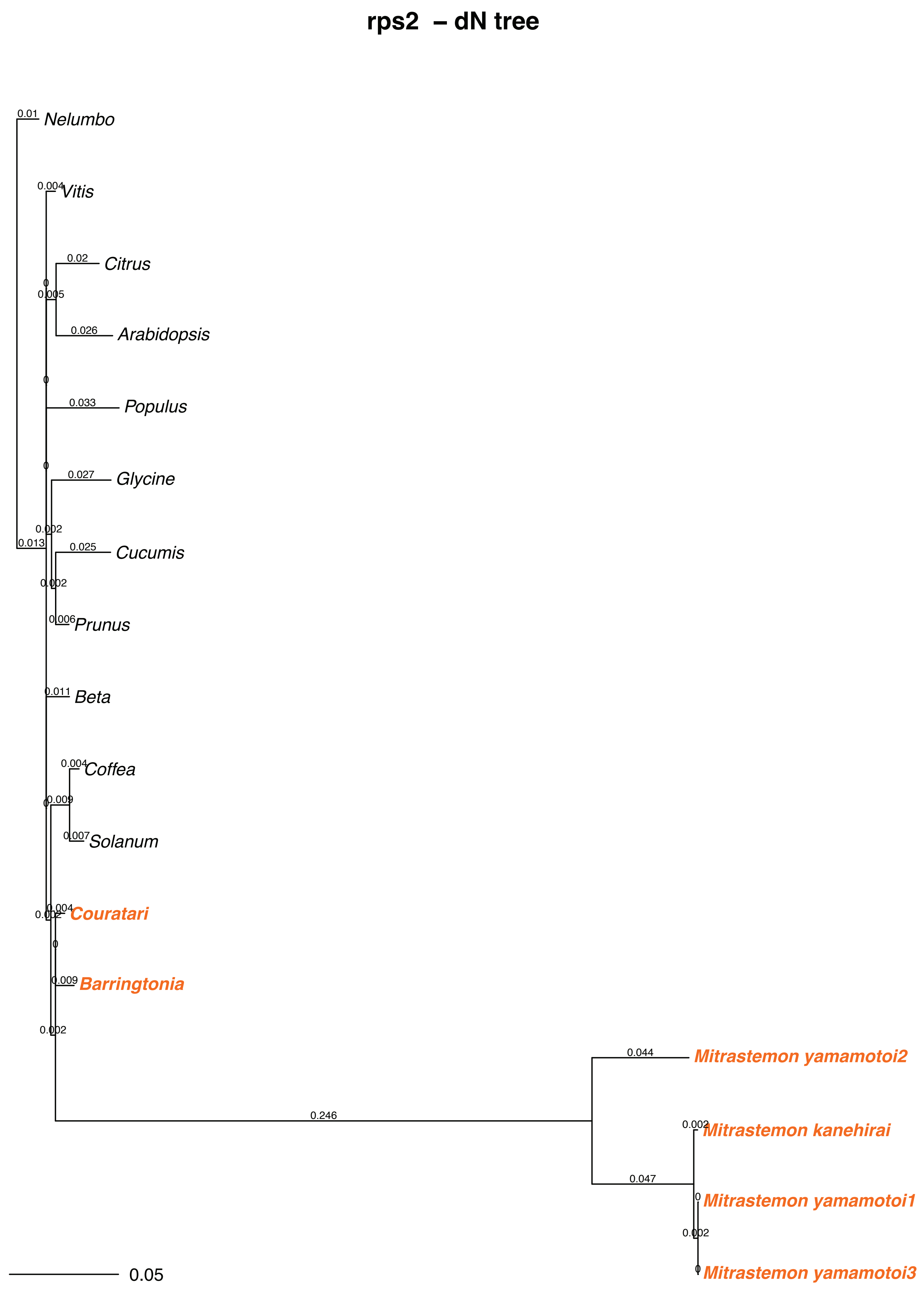

rps2 – dN tree
Nelumbo
Vitis
Citrus
Arabidopsis
Populus
Glycine
Cucumis
Prunus
Beta
Coffea
Solanum
Couratari
Barringtonia
Mitrastemon yamamotoi2
Mitrastemon kanehirai
Mitrastemon yamamotoi1
Mitrastemon yamamotoi3
0.01
0.004
0.02
0.005
0.026
0.033
0.027
0.002
0.013
0.025
0.006
0.011
0.009
0.007
0.044
0.246
0.047
0.05

### rps2 – dS tree

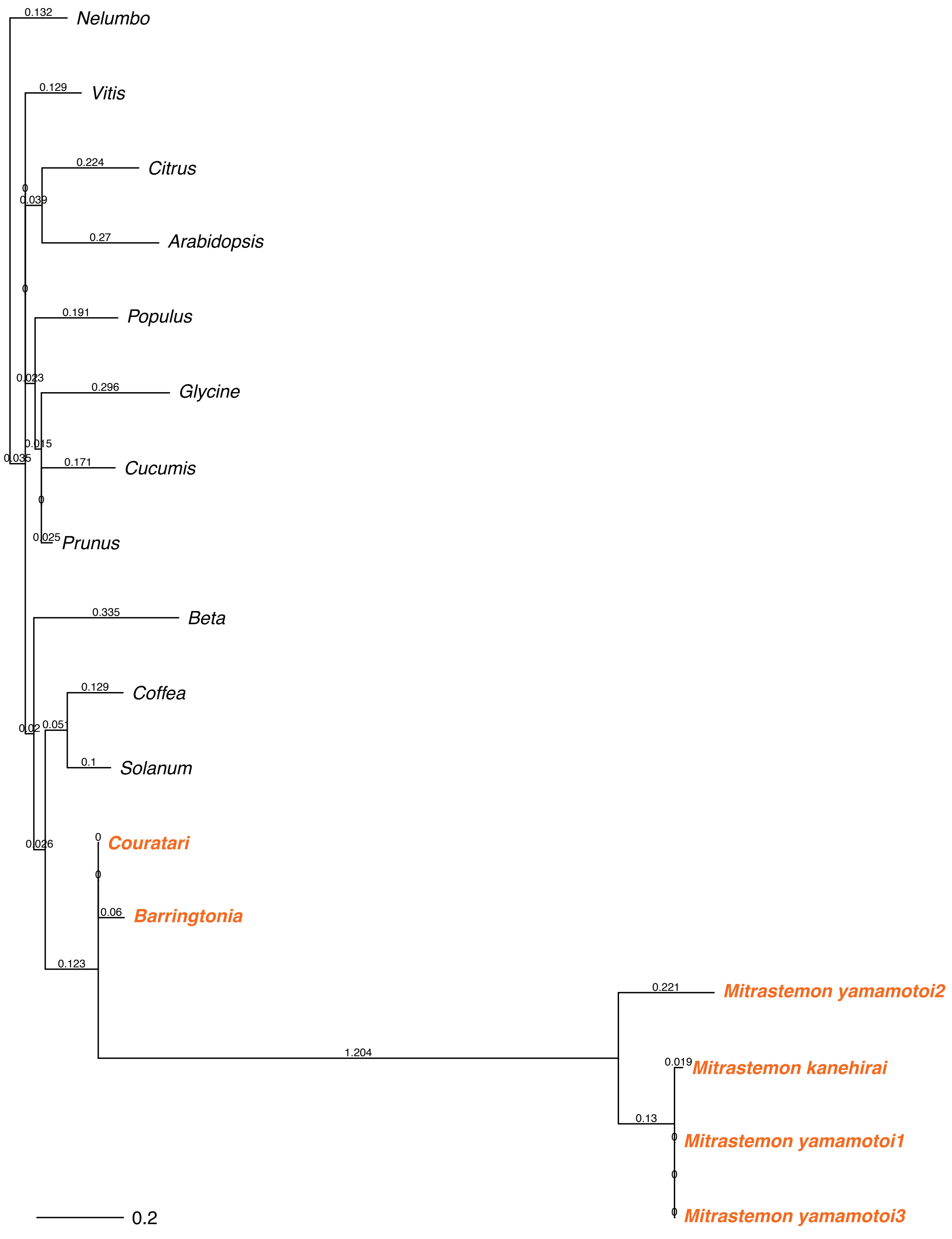
0.132
Nelumbo
0.129
Vitis
0.224
Citrus
0
0.039
0.27
Arabidopsis
0
0.191
Populus
0.023
0.296
Glycine
0.015
0.035
0.171
Cucumis
0
0.025
Prunus
0.335
Beta
0.129
Coffea
0.02
0.051
0.1
Solanum
0.026
0
Couratari
0
0.06
Barringtonia
0.123
0.221
Mitrastemon yamamotoi2
1.204
0.019
Mitrastemon kanehirai
0.13
0
Mitrastemon yamamotoi1
0
0
Mitrastemon yamamotoi3
0.2

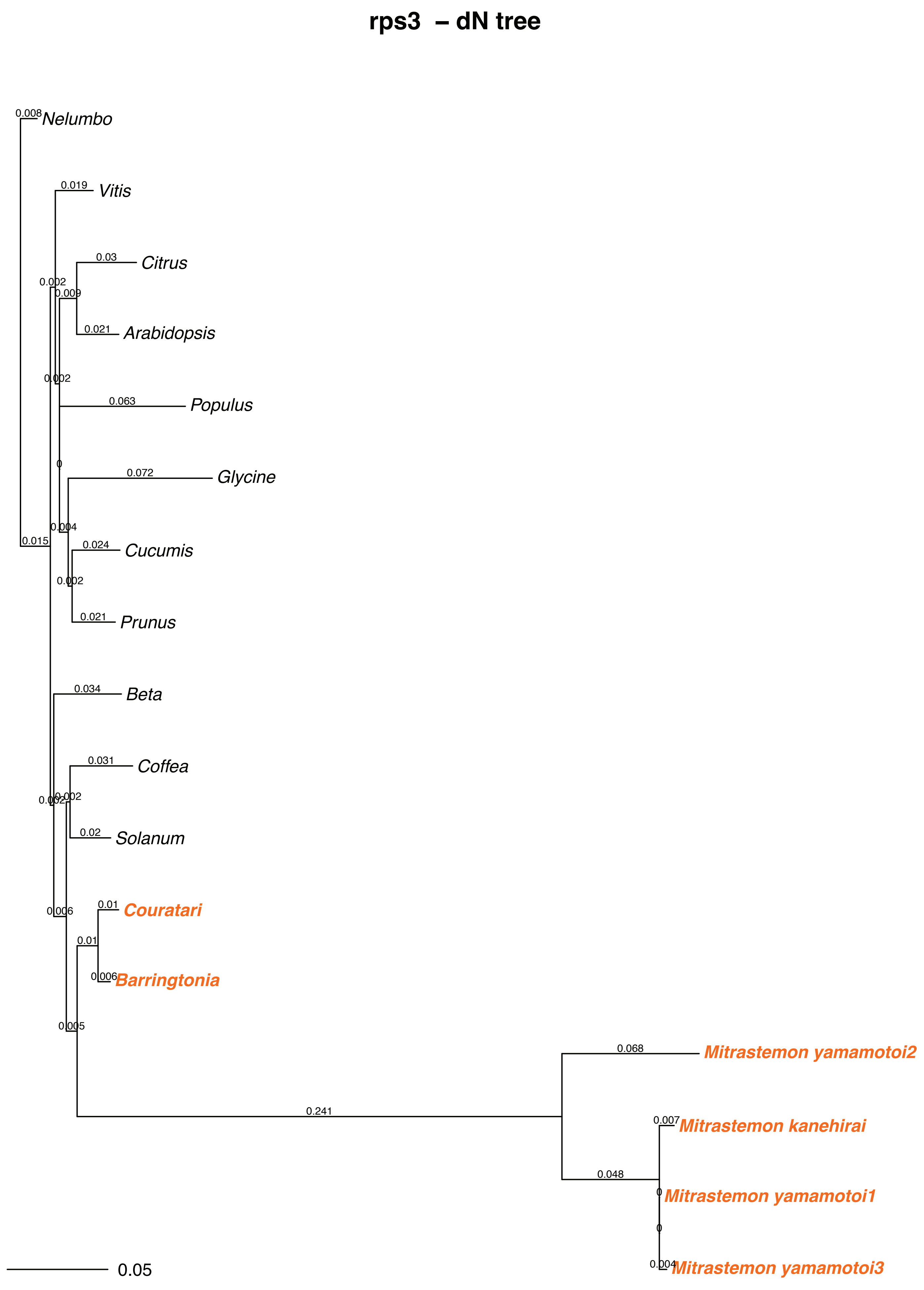

rps3 – dN tree
Nelumbo
Vitis
Citrus
Arabidopsis
Populus
Glycine
Cucumis
Prunus
Beta
Coffea
Solanum
Couratari
Barringtonia
Mitrastemon yamamotoi2
Mitrastemon kanehirai
Mitrastemon yamamotoi1
Mitrastemon yamamotoi3
0.008
0.019
0.03
0.002
0.009
0.021
0.002
0.063
0
0.072
0.004
0.015
0.024
0.002
0.021
0.034
0.031
0.002
0.002
0.02
0.01
0.006
0.01
0.006
0.005
0.068
0.241
0.007
0.048
0
0
0.004
0.05

#### rps3 – dS tree

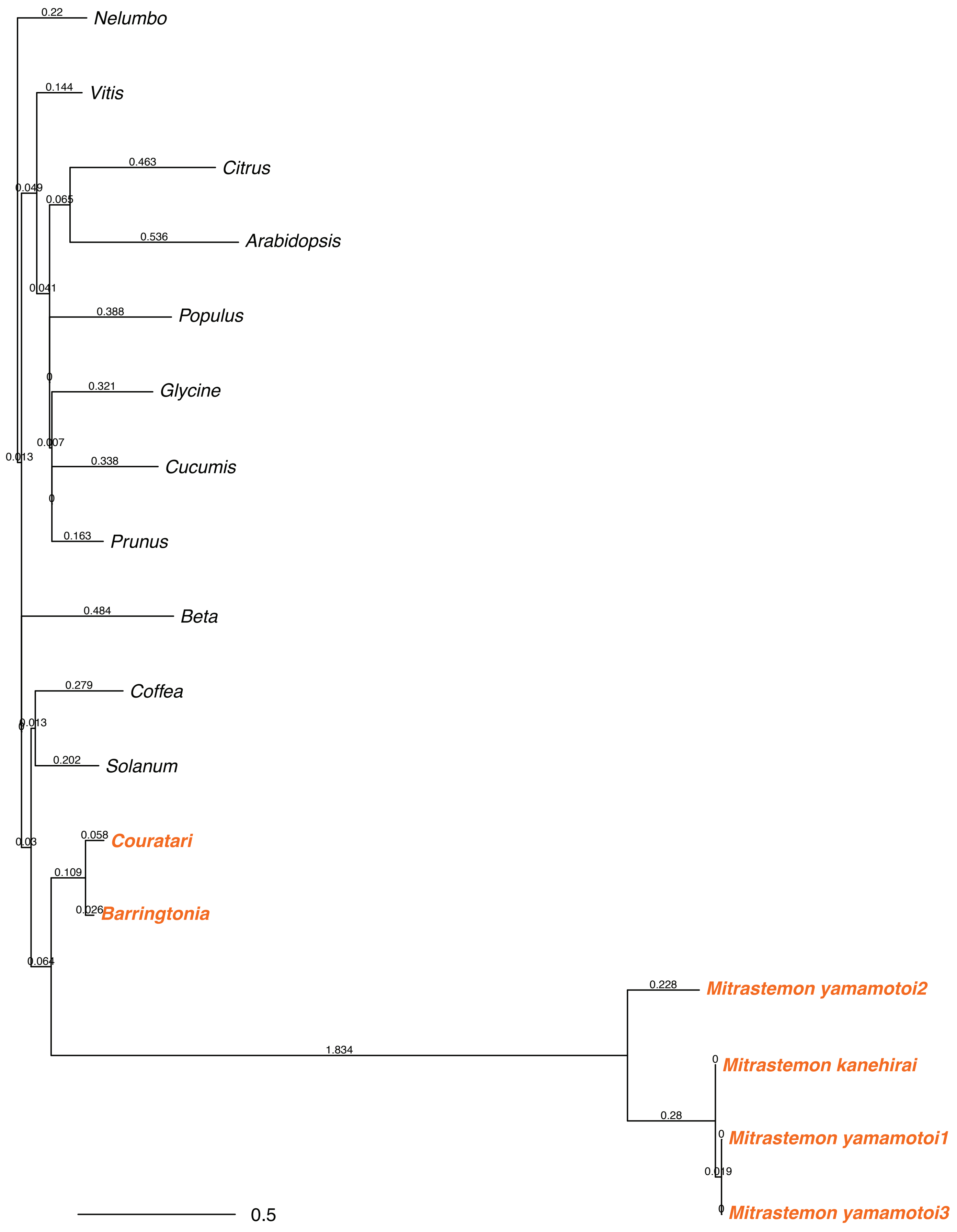
Nelumbo
0.22
Vitis
0.144
Citrus
0.463
Arabidopsis
0.536
0.049
0.065
0.041
Populus
0.388
0
Glycine
0.321
0.007
0.013
Cucumis
0.338
0
Prunus
0.163
Beta
0.484
Coffea
0.279
0.013
0
Solanum
0.202
Couratari
0.058
0.03
0.109
Barringtonia
0.026
0.064
Mitrastemon yamamotoi2
0.228
1.834
Mitrastemon kanehirai
0
0.28
Mitrastemon yamamotoi1
0
0.019
Mitrastemon yamamotoi3
0
0.5

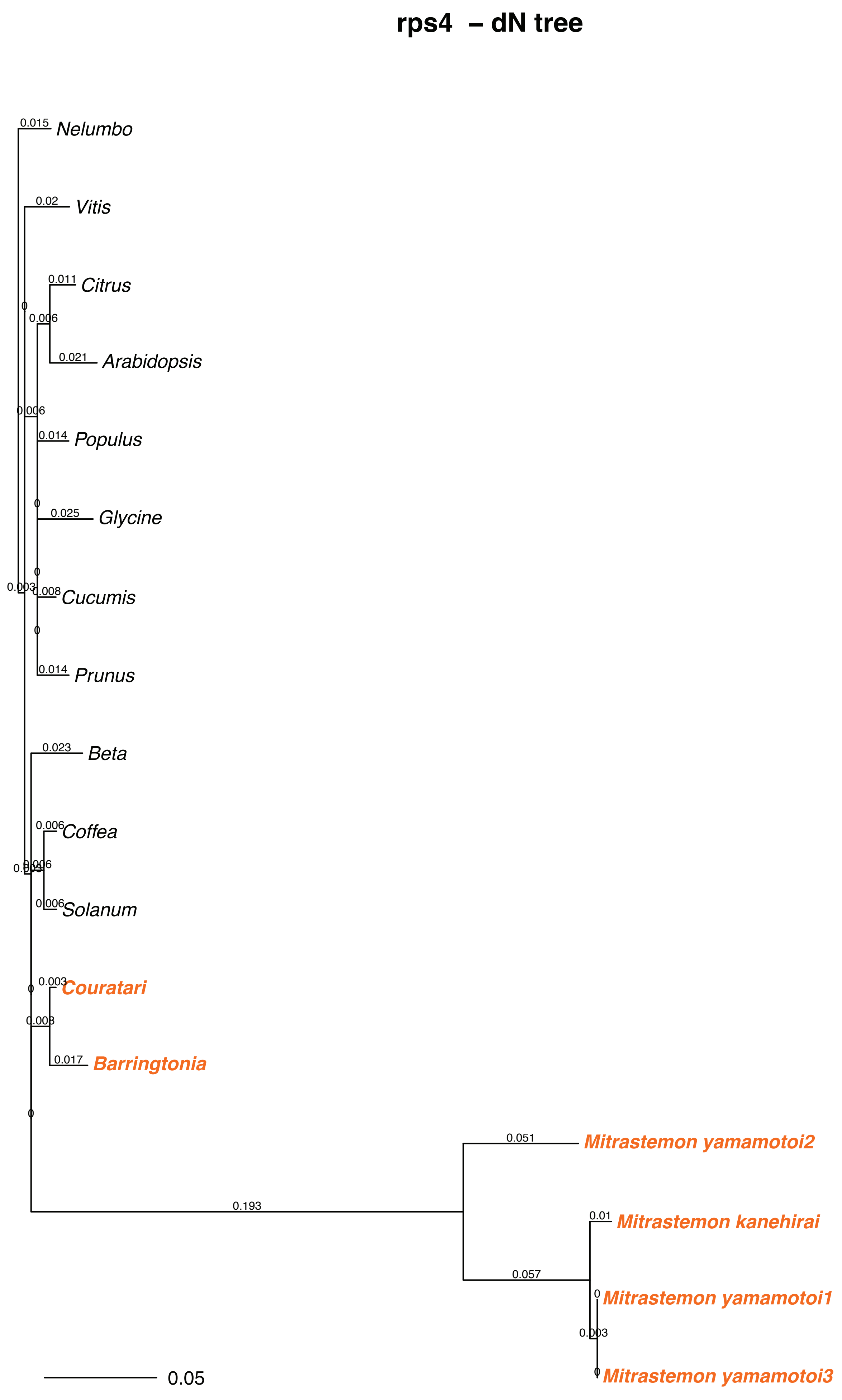
rps4 – dN tree
Nelumbo
Vitis
Citrus
Arabidopsis
Populus
Glycine
Cucumis
Prunus
Beta
Coffea
Solanum
Couratari
Barringtonia
Mitrastemon yamamotoi2
Mitrastemon kanehirai
Mitrastemon yamamotoi1
Mitrastemon yamamotoi3
0.015
0.02
0.011
0.021
0.006
0.014
0.025
0.008
0.003
0.023
0.003
0.017
0.051
0.193
0.01
0.057
0
0.05

#### rps4 – dS tree

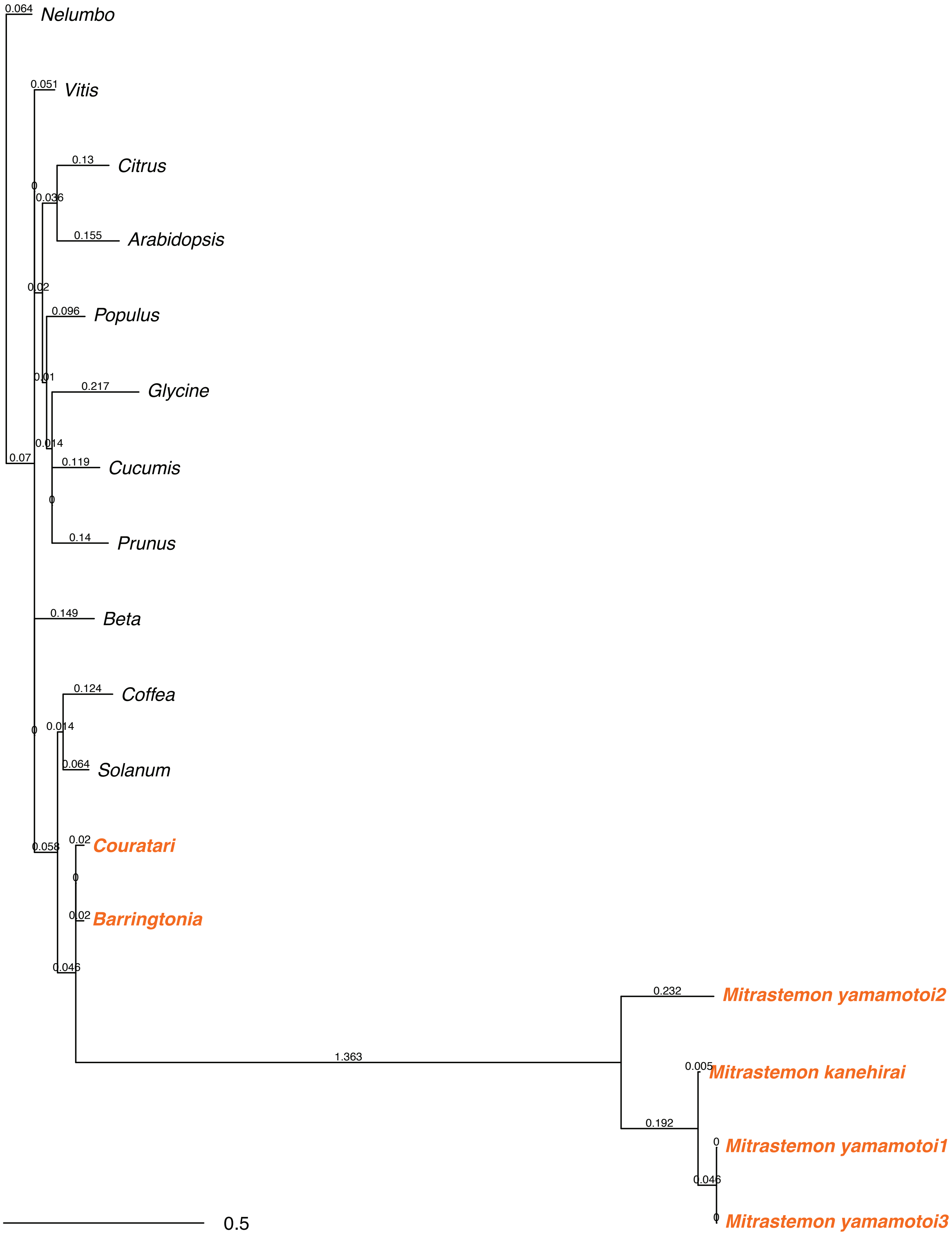

0.064
Nelumbo
0.051
Vitis
0.13
Citrus
0
0.036
0.155
Arabidopsis
0.02
0.096
Populus
0.01
0.217
Glycine
0.014
0.07
0.119
Cucumis
0
0.14
Prunus
0.149
Beta
0.124
Coffea
0
0.014
0.064
Solanum
0.058
0.02
Couratari
0
0.02
Barringtonia
0.046
0.232
Mitrastemon yamamotoi2
1.363
0.005
Mitrastemon kanehirai
0.192
0
Mitrastemon yamamotoi1
0.046
0
Mitrastemon yamamotoi3
0.5

#### rps7 – dN tree

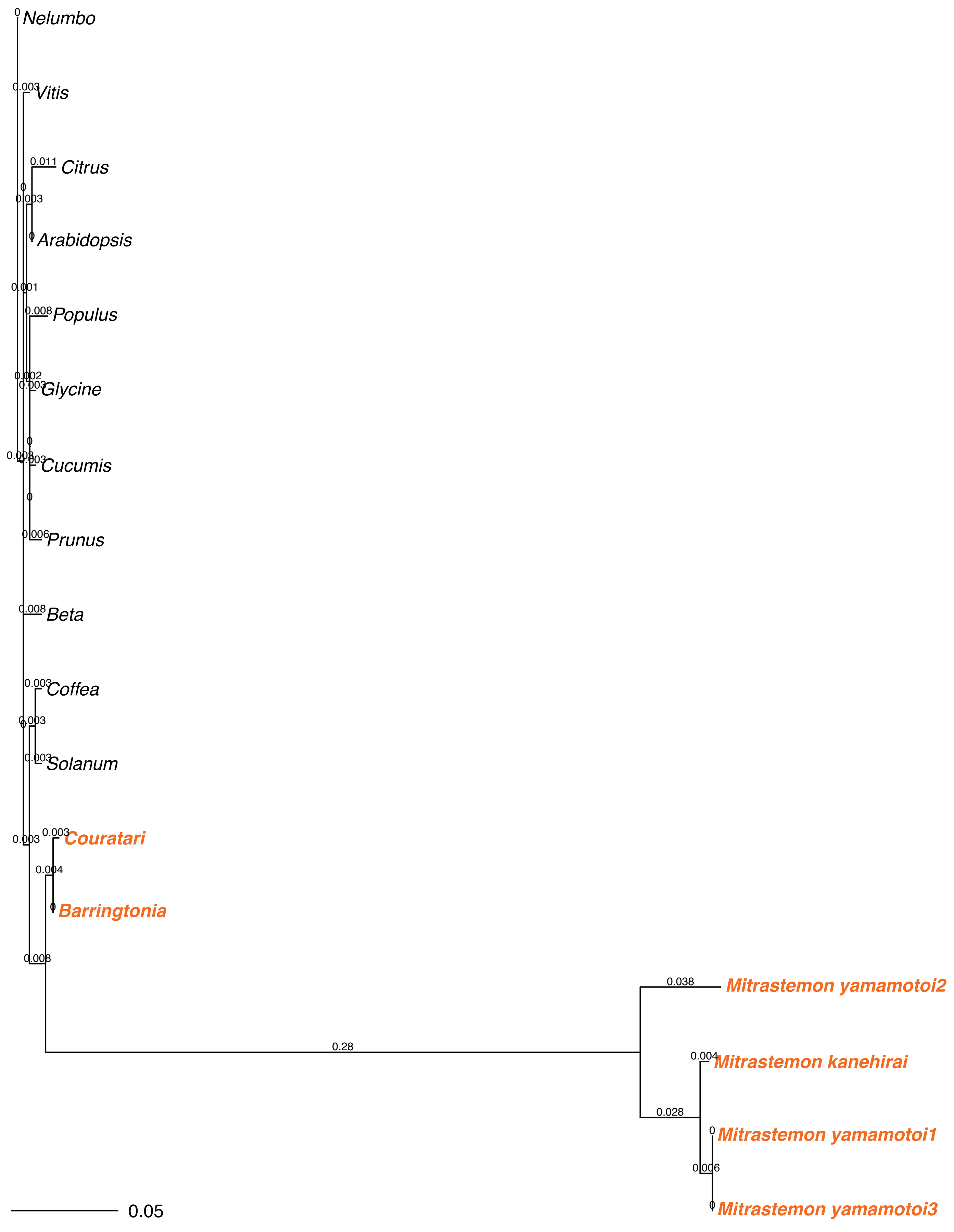

Nelumbo
Vitis
Citrus
Arabidopsis
Populus
Glycine
Cucumis
Prunus
Beta
Coffea
Solanum
Couratari
Barringtonia
Mitrastemon yamamotoi2
Mitrastemon kanehirai
Mitrastemon yamamotoi1
Mitrastemon yamamotoi3
0.28
0.038
0.028
0.05

### rps7 – dS tree

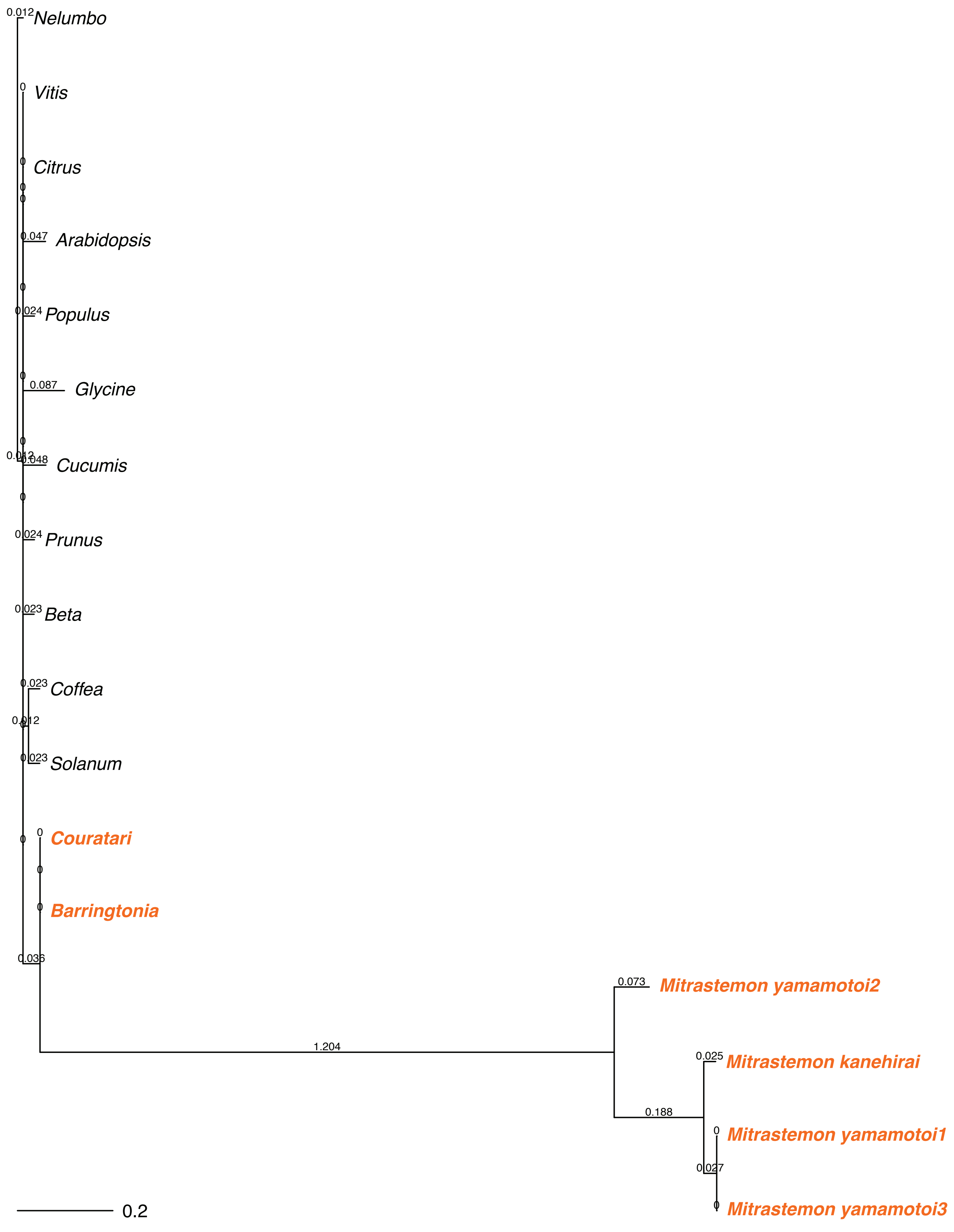

0.012 Nelumbo
0 Vitis
0 Citrus
0.047 Arabidopsis
0.024 Populus
0.087 Glycine
0.012
0.048 Cucumis
0.024 Prunus
0.023 Beta
0.023 Coffea
0.012
0.023 Solanum
0 Couratari
0 Barringtonia
0.036
1.204
0.073 Mitrastemon yamamotoi2
0.025 Mitrastemon kanehirai
0.188
0 Mitrastemon yamamotoi1
0.027
0 Mitrastemon yamamotoi3
0.2

#### rps8 – dN tree

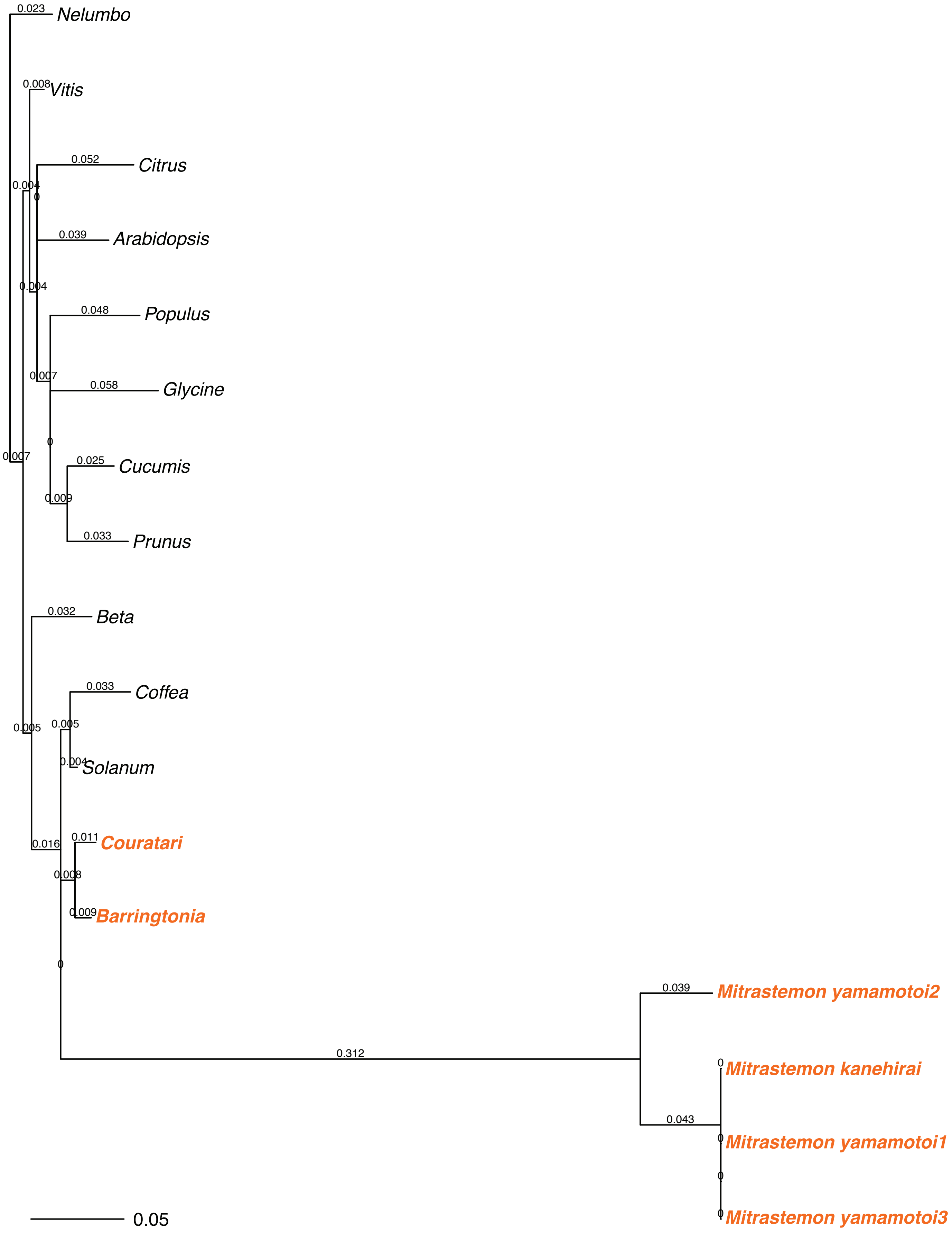

0.023 Nelumbo
0.008 Vitis
0.052 Citrus
0.004
0
0.039 Arabidopsis
0.004
0.048 Populus
0.007
0.058 Glycine
0
0.025 Cucumis
0.007
0.009
0.033 Prunus
0.032 Beta
0.033 Coffea
0.005
0.005
0.004 Solanum
0.011 Couratari
0.016
0.008
0.009 Barringtonia
0
0.039 Mitrastemon yamamotoi2
0.312
0 Mitrastemon kanehirai
0.043
0 Mitrastemon yamamotoi1
0
0 Mitrastemon yamamotoi3
0.05

#### rps8 – dS tree

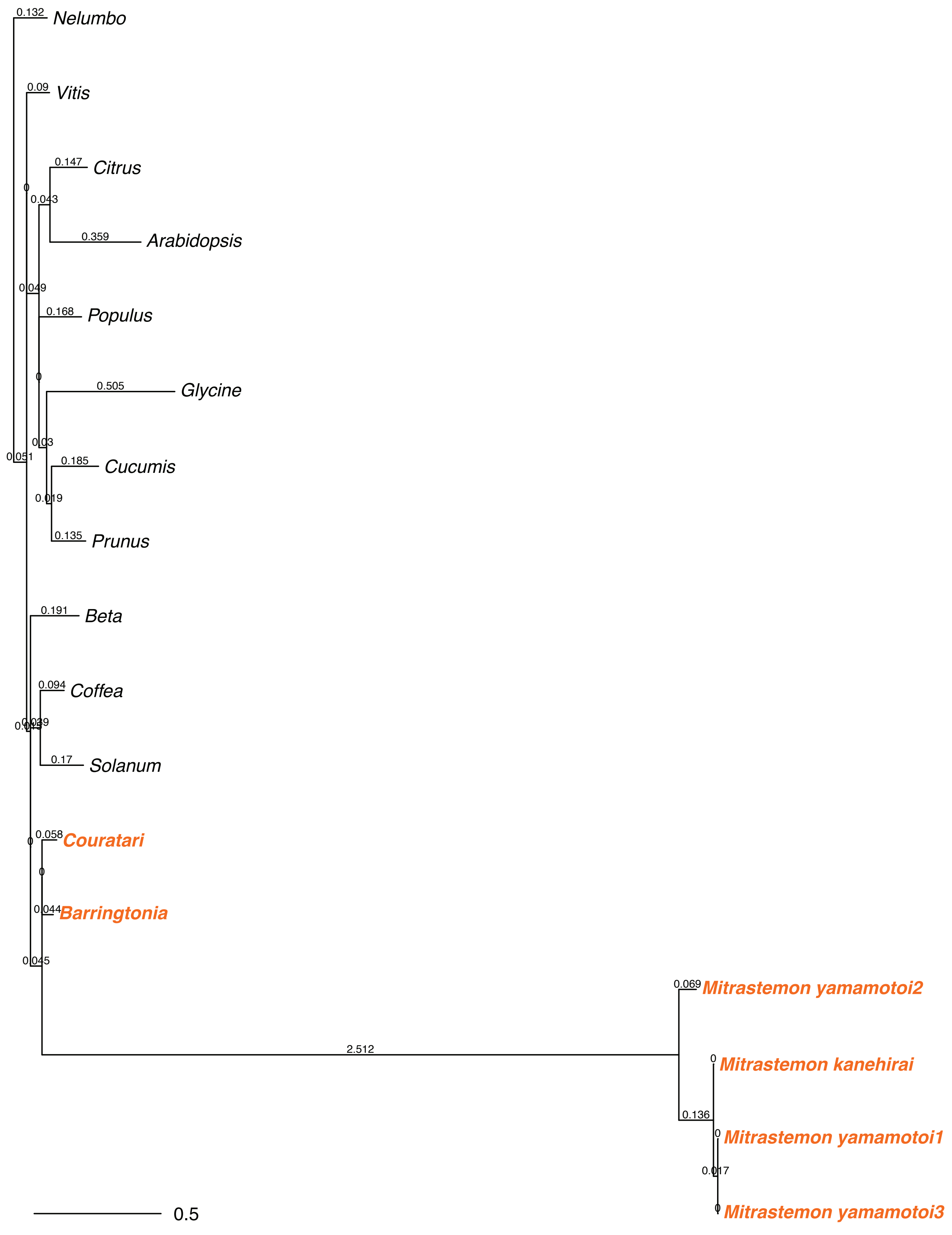

Nelumbo
Vitis
Citrus
Arabidopsis
Populus
Glycine
Cucumis
Prunus
Beta
Coffea
Solanum
Couratari
Barringtonia
Mitrastemon yamamotoi2
Mitrastemon kanehirai
Mitrastemon yamamotoi1
Mitrastemon yamamotoi3
0.132
0.09
0.147
0.359
0.043
0.049
0.168
0.505
0.03
0.185
0.019
0.135
0.051
0.191
0.094
0.039
0.17
0.058
0.044
0.045
2.512
0.069
0.136
0.017
0.5

#### rps11 – dN tree

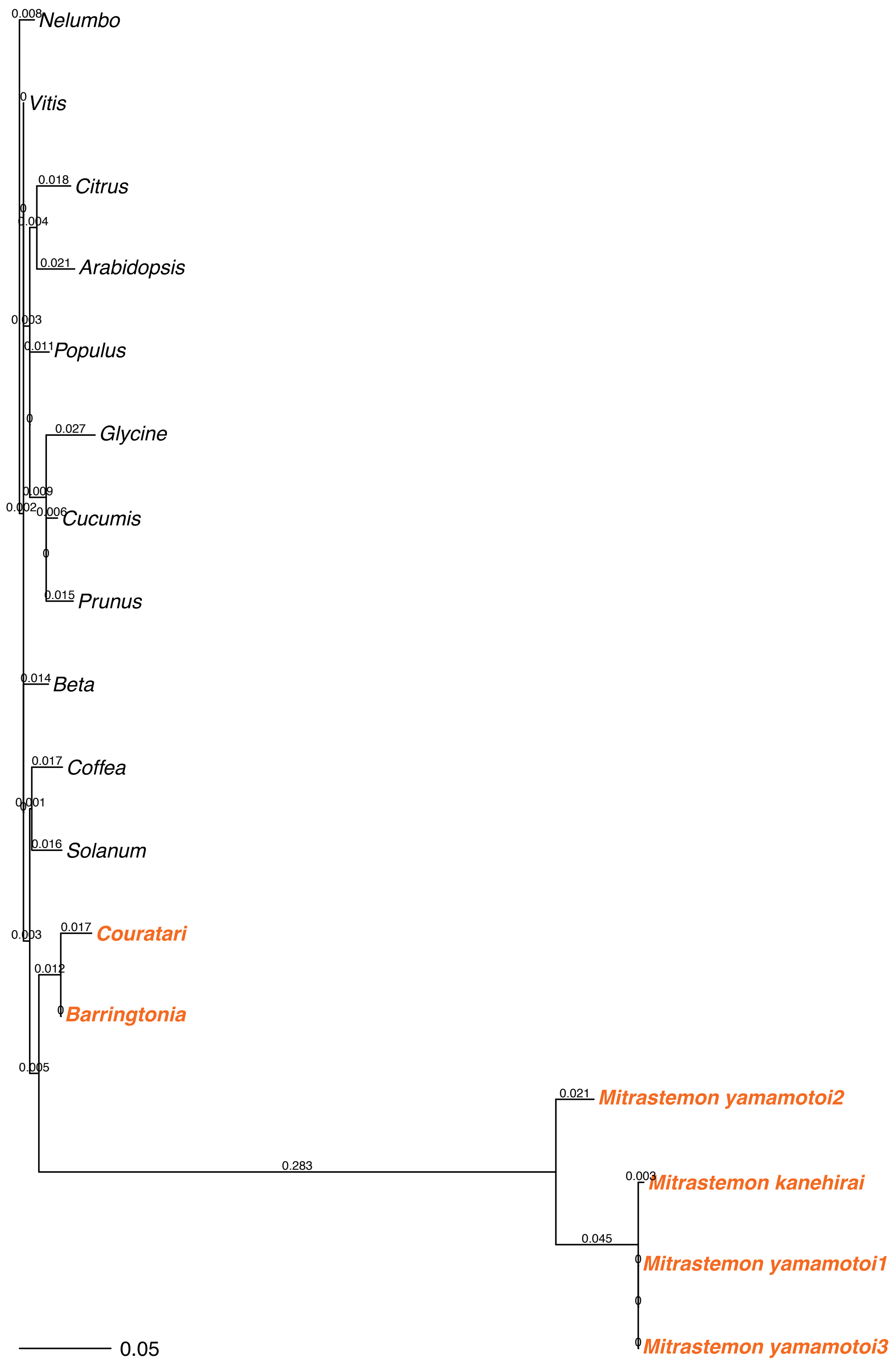

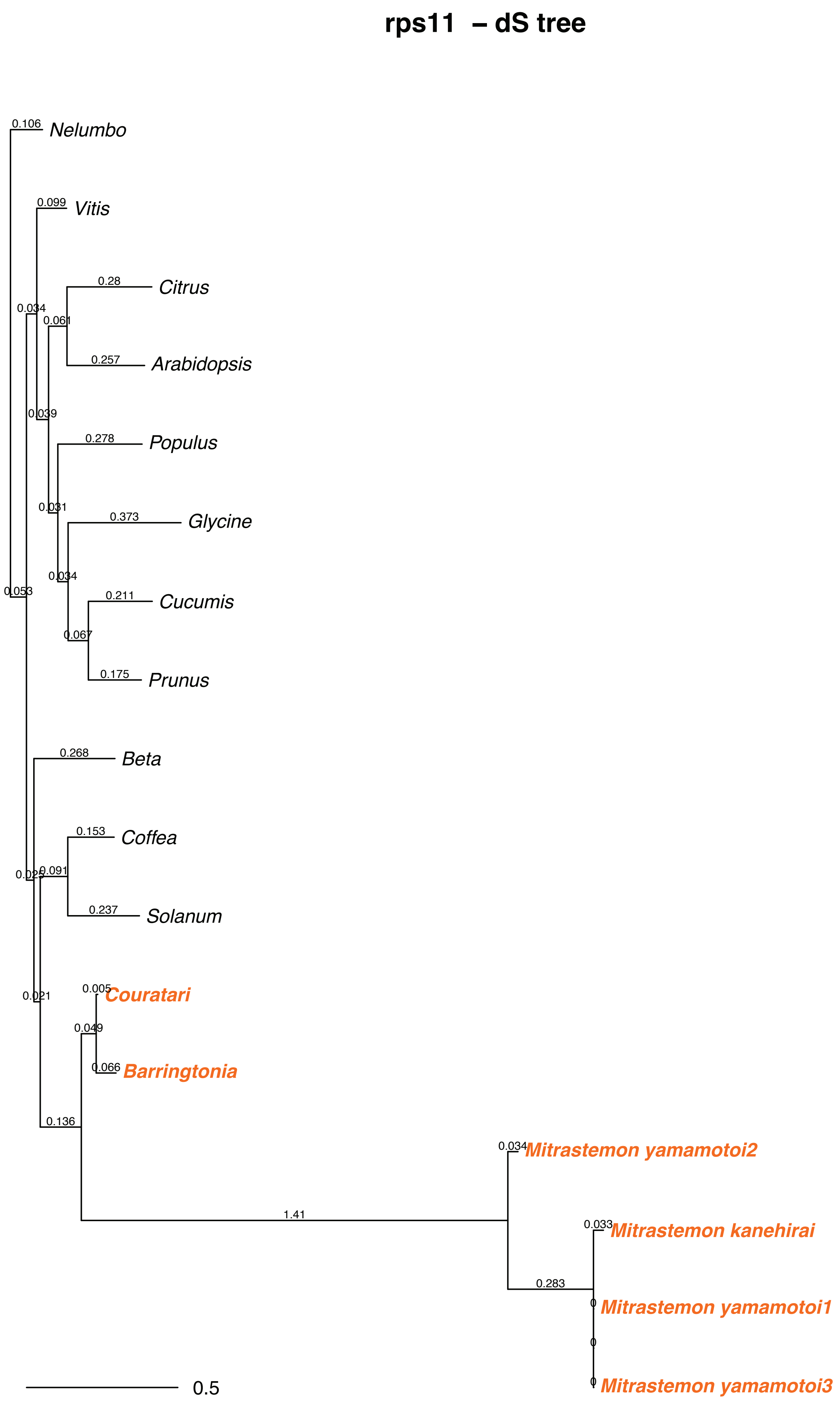
rps11 – dS tree
Nelumbo
Vitis
Citrus
Arabidopsis
Populus
Glycine
Cucumis
Prunus
Beta
Coffea
Solanum
Couratari
Barringtonia
Mitrastemon yamamotoi2
Mitrastemon kanehirai
Mitrastemon yamamotoi1
Mitrastemon yamamotoi3
0.106
0.099
0.28
0.257
0.034
0.061
0.039
0.278
0.031
0.373
0.034
0.211
0.067
0.175
0.053
0.268
0.153
0.091
0.237
0.021
0.005
0.049
0.066
0.136
1.41
0.034
0.033
0.283
0
0
0
0.5

#### rps12 – dN tree

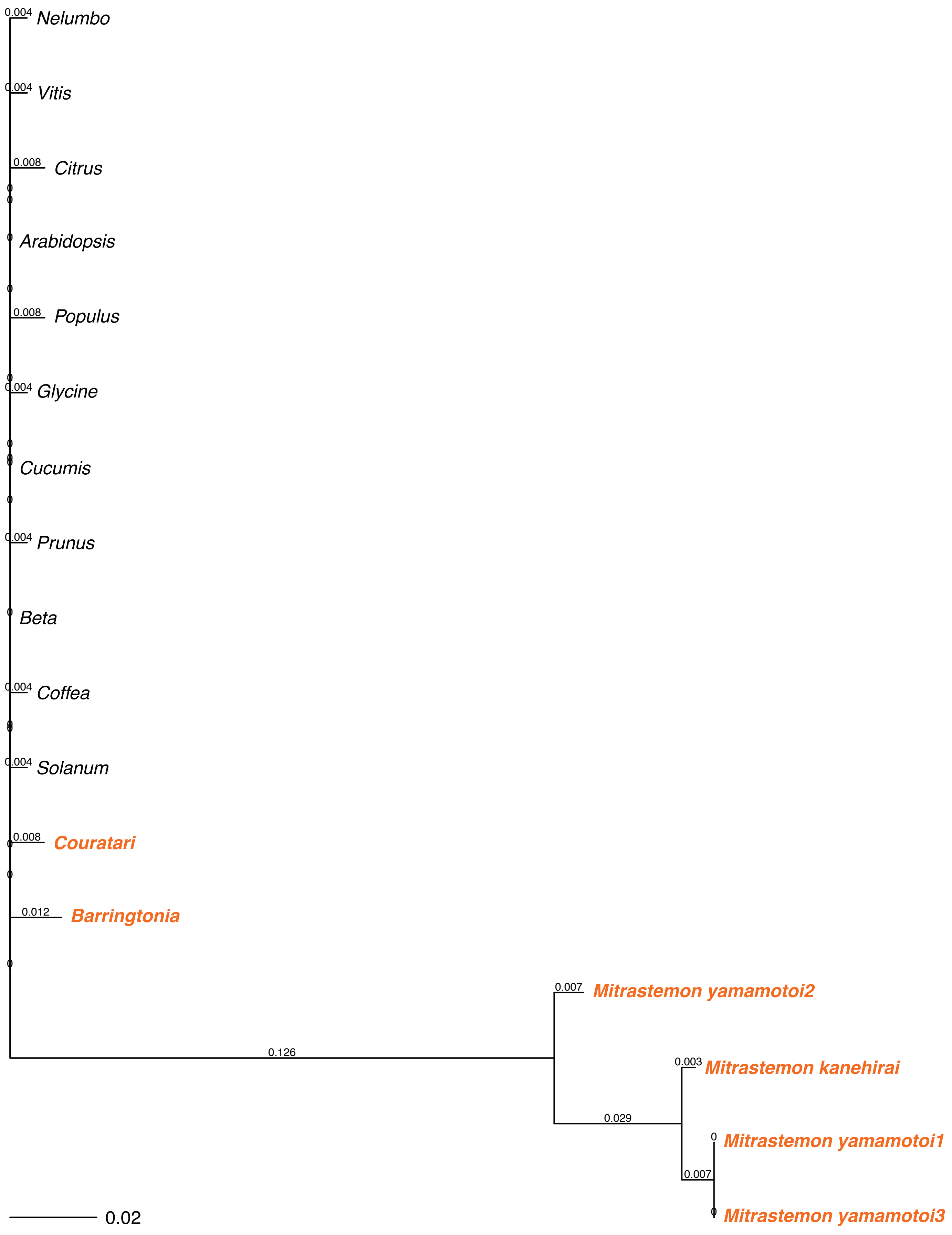

### rps12 – dS tree

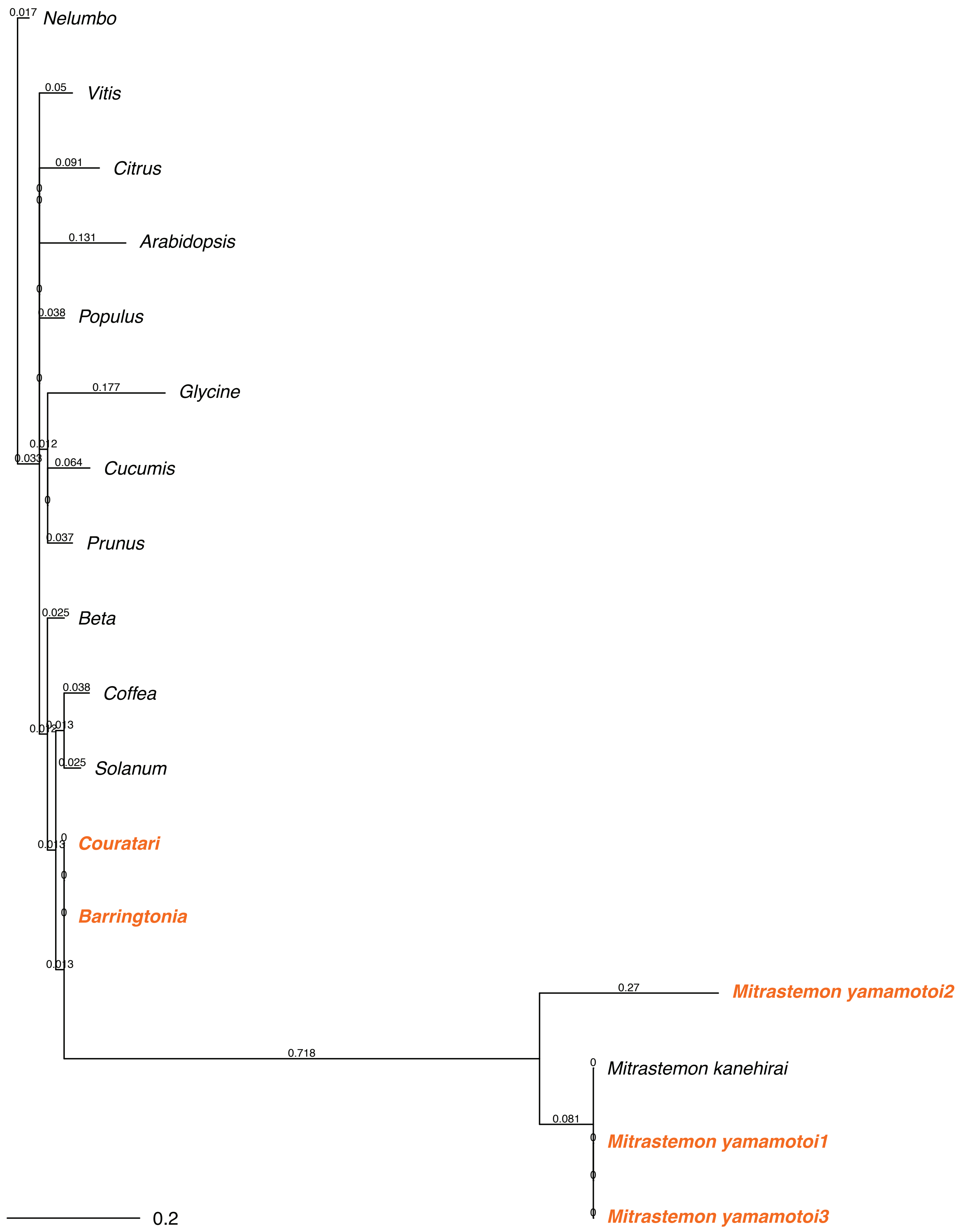

#### rps14 – dN tree

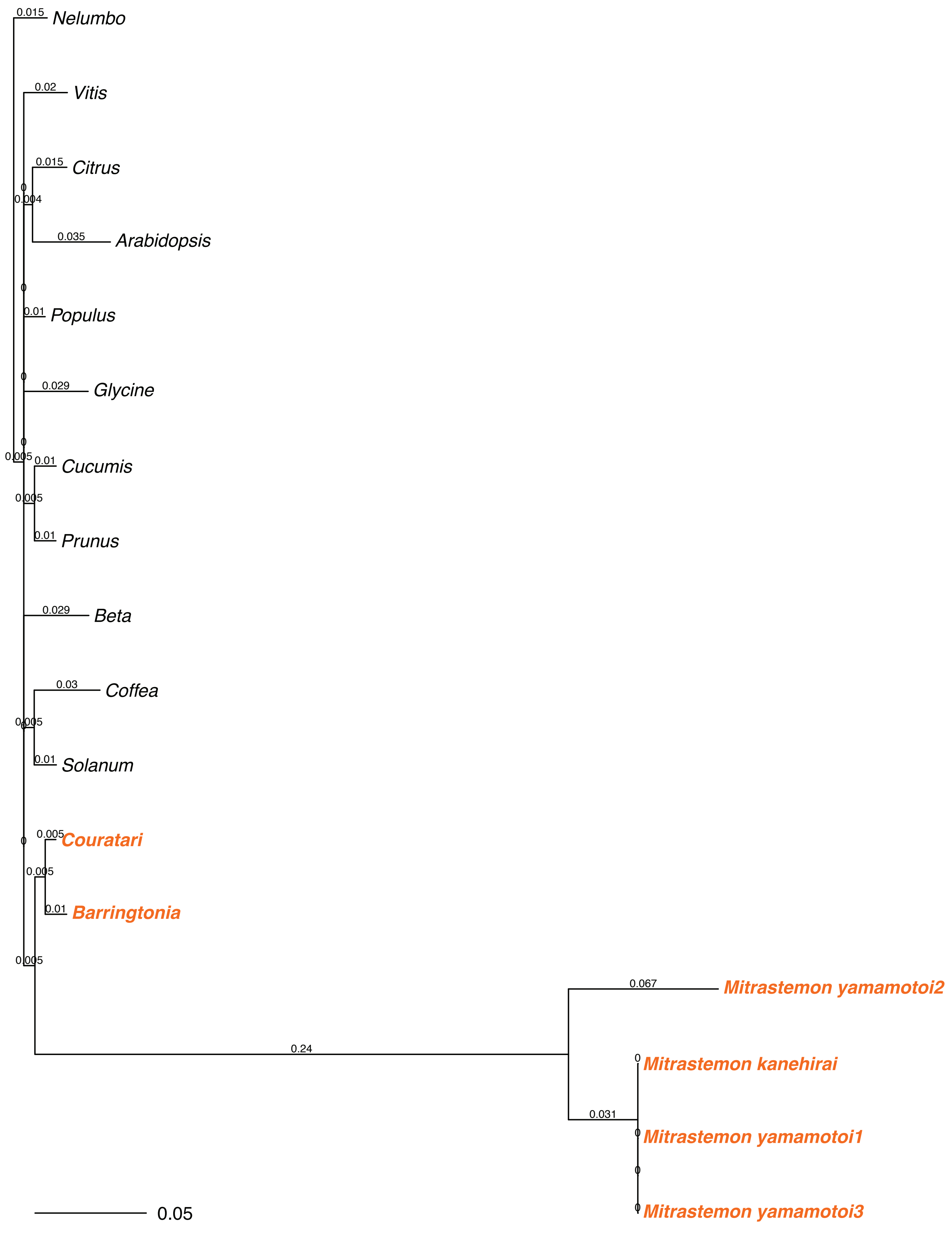

#### rps14 – dS tree

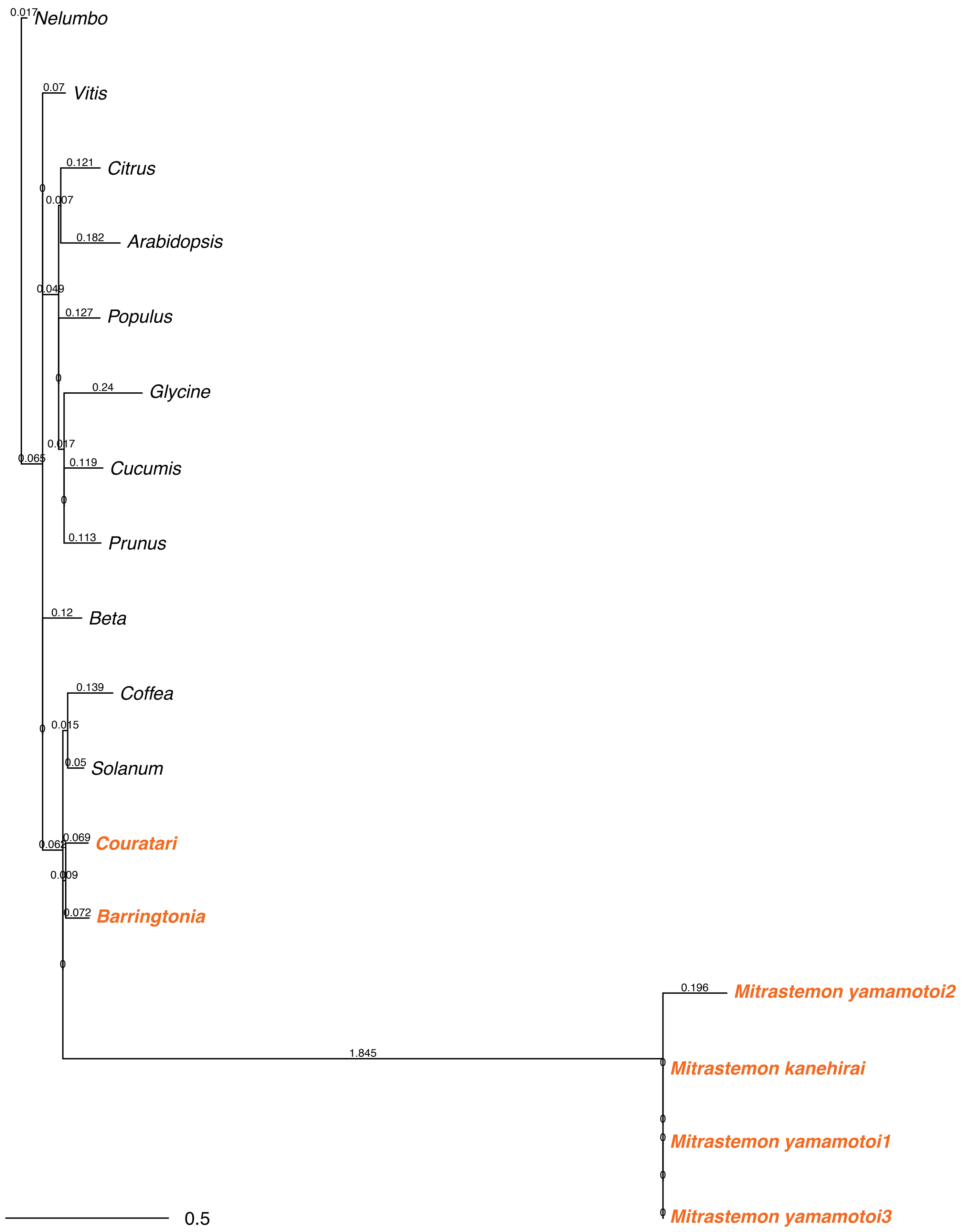

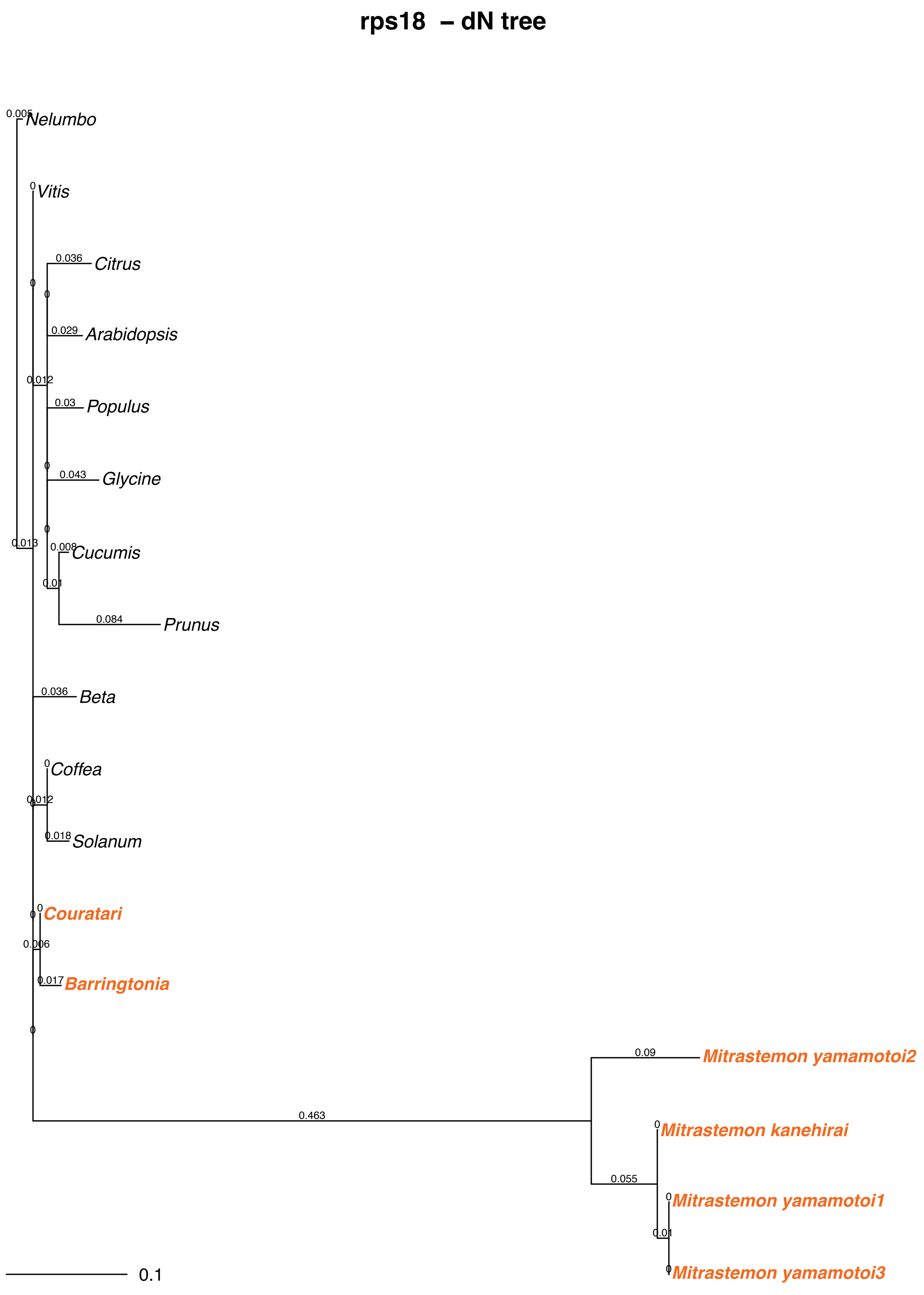
rps18 – dN tree
Nelumbo
Vitis
Citrus
Arabidopsis
Populus
Glycine
Cucumis
Prunus
Beta
Coffea
Solanum
Couratari
Barringtonia
Mitrastemon yamamotoi2
Mitrastemon kanehirai
Mitrastemon yamamotoi1
Mitrastemon yamamotoi3
0.005
0.036
0.029
0.012
0.03
0.043
0.013
0.008
0.01
0.084
0.018
0.006
0.017
0.09
0.463
0.055
0.1

#### rps18 – dS tree

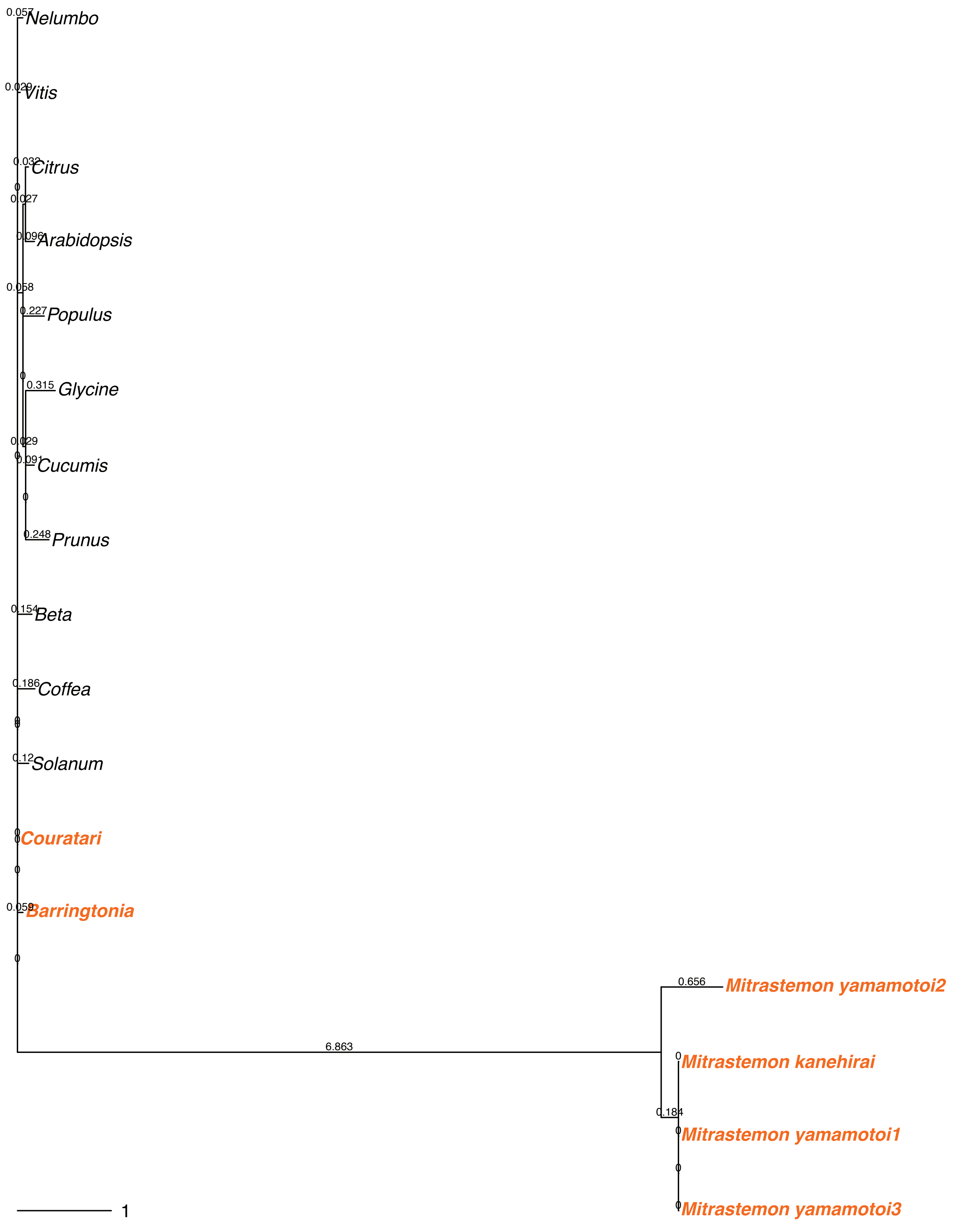

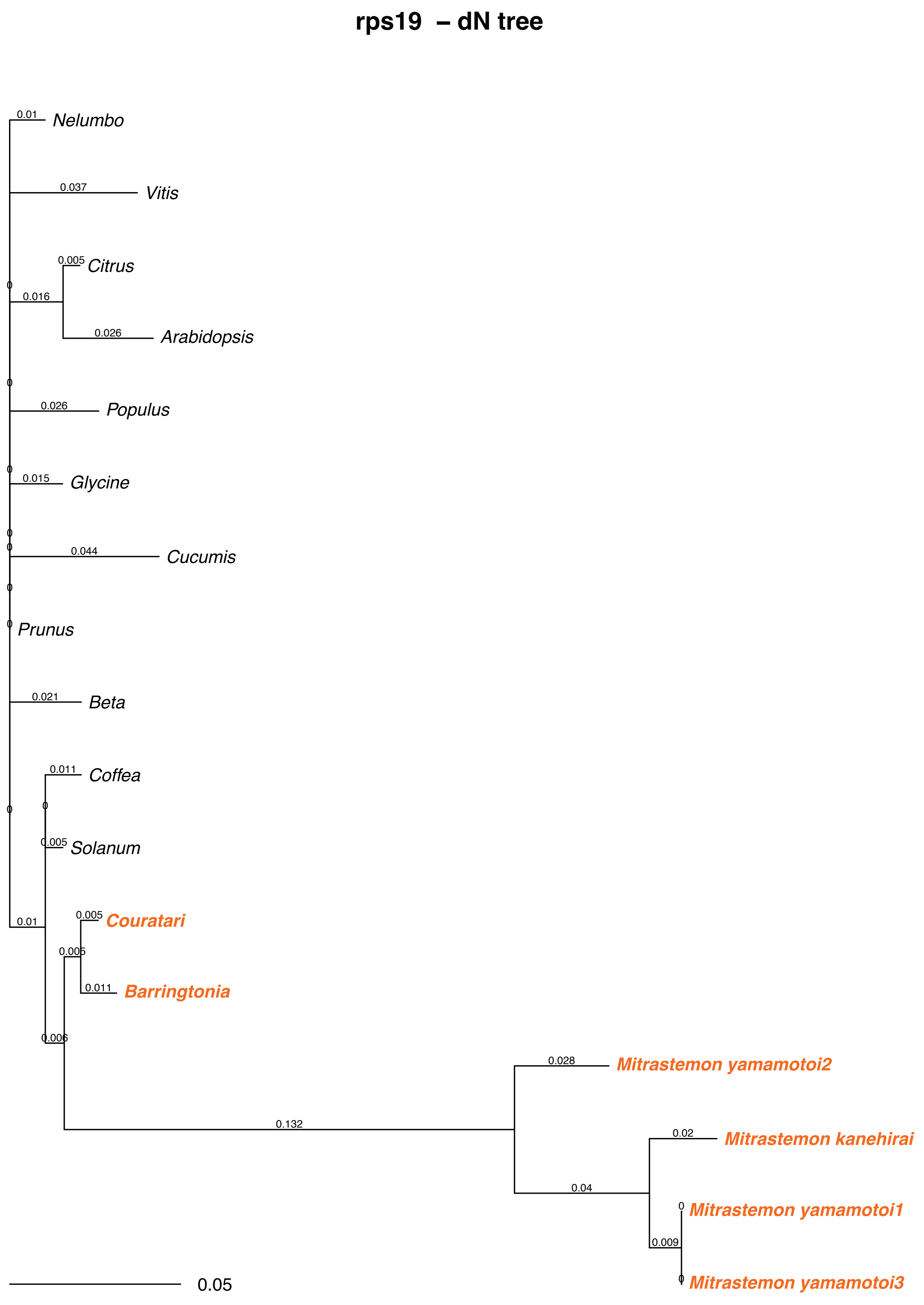
rps19 – dN tree
0.01
Nelumbo
0.037
Vitis
0.005
Citrus
0.016
0.026
Arabidopsis
0.026
Populus
0.015
Glycine
0.044
Cucumis
Prunus
0.021
Beta
0.011
Coffea
0.005
Solanum
0.005
Couratari
0.01
0.005
0.011
Barringtonia
0.006
0.028
Mitrastemon yamamotoi2
0.132
0.02
Mitrastemon kanehirai
0.04
Mitrastemon yamamotoi1
0.009
Mitrastemon yamamotoi3
0.05

#### rps19 – dS tree

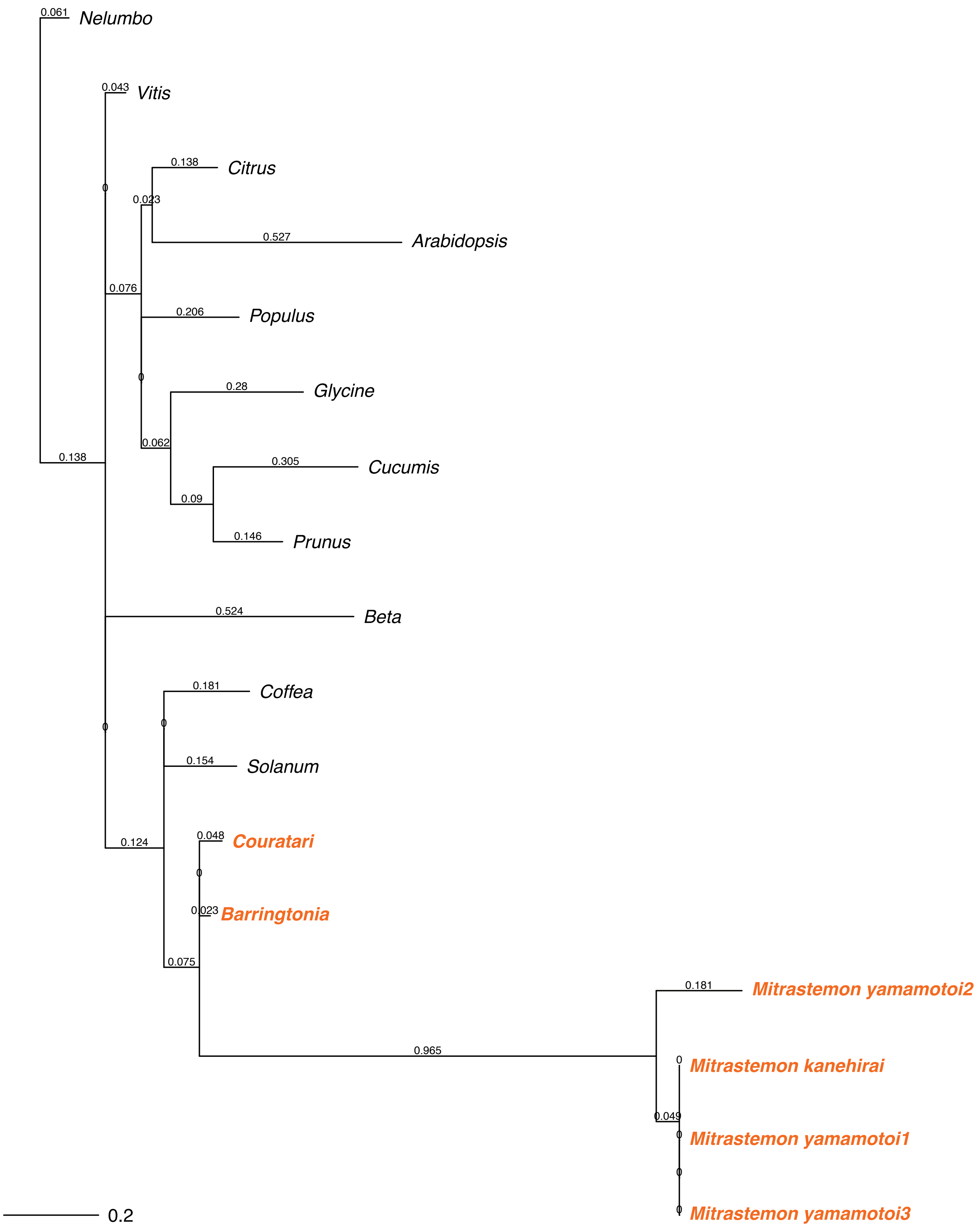
